# Recombination and the role of pseudo-overdominance in polyploid evolution

**DOI:** 10.1101/2025.02.28.640841

**Authors:** William W. Booker, Daniel R. Schrider

## Abstract

Natural selection is an imperfect force that can under some conditions fail to prevent the buildup of deleterious mutations. Small population sizes and the lack of recombination are two such scenarios that reduce the efficiency of selection. Under these conditions, the disconnect between deleterious genetic load and individual fitness due to the masking of recessive deleterious mutations in heterozygous individuals may result in the emergence of pseudo-overdominance, wherein the buildup of haplotypes with complementary sets of deleterious mutations results in apparent heterozygote advantage and an increase in linked neutral diversity. In polyploids, the presence of additional allelic copies magnifies this masking effect and may therefore increase the probability of pseudo-overdominance. Here, we simulate the evolution of small diploid and autotetraploid populations to identify the conditions that support the evolution of pseudo-overdominance. We discover that pseudo-overdominance evolves under a much wider range of parameters in autotetraploids than in diploids with identical population sizes, and that in many parts of parameter space there is an inverse relationship between fitness and recombination rate. These results imply that pseudo-overdominance may be more common than previously thought. We conclude by discussing the current evidence for pseudo-overdominance in species with polyploid histories, as well as its implications in agriculture due to the prevalence of polyploidy in crops.

**ARTICLE SUMMARY:** Pseudo-overdominance is an extreme evolutionary phenomenon in which low recombination rates result in the buildup of alternative mutations on disparate haplotypes, rendering recombination events maladaptive because they disrupt the masking of recessive deleterious alleles. In polyploids, additional chromosomes accentuate this masking effect, potentially increasing the likelihood of pseudo-overdominance. We used forward-in-time simulations to test the hypothesis that polyploids are more likely to experience pseudo-overdominance–and find strong evidence in support of this hypothesis. Our results have implications for the evolution of recombination rates in all species, the evolution of polyploids generally, and the efficacy of selection in polyploid crops.

## INTRODUCTION

Recombination is a fundamental process in shaping the evolution of species and their genomes. In its absence, mutations accumulate on chromosomes without the ability to be purged outside of the destruction of the entire chromosome’s lineage (Muller 1964; Felsenstein 1974). When present, recombination breaks down linkage disequilibrium and new favorable mutations can find their way onto common alleles while deleterious mutations can be removed from those backgrounds, enabling a more efficient traversal to the peaks of the fitness landscape (Crow and Kimura 1965; Hill and Robertson 1966; Felsenstein 1974; Barton 1995). One might therefore expect selection to drive recombination rates higher and higher. However, plants and animals are generally limited to having roughly two crossovers per generation per chromosome—albeit with substantial variation across species (Wilfert et al. 2007; Stapley et al. 2017; Brazier and Glémin 2022). This phenomenon is in part due to molecular constraints such as crossover interference (Muller 1916; Wang et al. 2015; Ritz et al. 2017), but may also result from the deleterious effects of too much recombination: recombination events can break up haplotypes that have accumulated multiple beneficial alleles (Charlesworth and Charlesworth 1975; Barton and Charlesworth 1998) termed ‘recombination load’, (Charlesworth and Charlesworth 1975; Barton and Charlesworth 1998).

Pseudo-overdominance presents an extreme evolutionary phenomenon in which evolving higher rates of recombination is maladaptive (Frydenberg 1963; Sved 1968; Ohta and Kimura 1970; Ohta 1971; Sved 1971; Sved 1972; Ohta 1973; Charlesworth 1991; Becher et al. 2020; Waller 2021). Pseudo-overdominance occurs when extremely low recombination rates result in the buildup of deleterious mutations on disparate haplotypic backgrounds causing heterozygotes with alternate high-load haplotypes to be the most fit. Because mutations are unlikely to occur independently at the same site, low recombination regions of the genome can accumulate high levels of mutational load when mutations are at least partially recessive and therefore the load is only expressed in the homozygous state. This will result in an excess of genetic diversity relative to the neutral expectation (Pamilo and Pálsson 1998), not only for the deleterious mutations themselves but also for linked neutral alleles (with the latter effect said to be a consequence of associative overdominance; Frydenberg 1963). When pseudo-overdominant dynamics are operating, selection will disfavor mutations that increase recombination rates (Pálsson 2002), suggesting that selection could also favor modifiers that decrease recombination under pseudo-overdominance.

Pseudo- and associative overdominance have received relatively little attention, but signatures of these dynamics have been detected in *Drosophila* (Latter 1998; Becher et al. 2020) and humans (Gilbert et al. 2020), and have been suggested as a mechanism shaping the heterogeneous density of genetic diversity and load across the genomes of additional plant and animal species (Waller 2021). Although theory suggests these dynamics are most likely to occur in smaller, inbred populations (Zhao and Charlesworth 2016), they may also occur in larger populations under some conditions. For example, population bottlenecks can push populations with larger census sizes into a pseudo-overdominant state (Bierne et al. 2000). Moreover, even in large populations that have not experienced bottlenecks the reduced *N_e_* (effective population size) of low recombination regions caused by background selection and selective interference with other loci under selection could also subject these regions to the conditions in which pseudo-overdominance dynamics can occur (Zhao and Charlesworth 2016; Charlesworth and Jensen 2021).

The equilibrium state of a finite diploid population under pseudo-overdominant dynamics is that of multiple haplotypes with complementary deleterious mutations segregating in the population at frequencies relative to *s* (the selection coefficient) after alternative haplotypes have been lost to drift (Gilbert et al. 2020). Effectively, this means that although individual fitness is high, roughly half of all offspring are homozygous for a high-load haplotype. However, although rarely considered, diploidy is not the sole ploidal state in biology, and tetraploids under the same equilibrium dynamics would produce between 0-25% homozygote high-load offspring depending on the allelic state of the parents (with the maximum fraction of high-load offspring produced by mating between two AAAa parents, where A and a represent the two haplotypes). Higher-level ploidies would produce even fewer high-load offspring.

Polyploidization has shaped the evolutionary history of nearly all branches of the tree of life (Otto and Whitton 2000; Gregory and Mable 2005; Breuert et al. 2006; Wood et al. 2009; Albertin and Marullo 2012; Barker et al. 2016; Zhan et al. 2016; Van de Peer et al. 2017; Li et al. 2018; One Thousand Plant Transcriptomes Initiative 2019). Although there is considerable debate over any paradigmatic evolutionary trajectory of polyploids, several aspects of polyploid formation may promote pseudo-overdominant dynamics in polyploids specifically. First, due to the nature of polyploidization, all newly formed polyploids go through an initial population bottleneck significantly lowering their effective population size (Stebbins 1950). Second, the multiplication of chromosomes in a cell causes widespread genomic instability presenting several challenges to undergoing regular meiosis, and evidence suggests there exists a strong pressure to evolve lower recombination rates in newly formed polyploids (Morrison and Rajhathy 1960; Reddi 1970; Wu et al. 2014; Bomblies et al. 2016; Morgan et al. 2021)–often below that of their diploid progenitors (Mulligan 1967; Yant et al. 2013). Combined with the relative rarity of homozygotes in polyploids compared to diploids at identical allele frequencies and the corresponding greater masking of deleterious alleles, these observations suggest pseudo-overdominance may occur more often in polyploids as compared to other taxa.

In this study, we investigate the conditions in which pseudo-overdominant dynamics are present in autotetraploids. Using forward-in-time simulations, we analyze the relationship between fitness and demography as a function of ploidy, population size, mutation rate, recombination rate, and the strength of selection. We conduct simulations under a wide variety of parameters, attempting to identify the parameter space where pseudo-overdominant dynamics are favored in diploid and autotetraploid populations. Additionally, we describe the complex relationship between fitness and recombination rate in small populations, as well as the long-term trajectory of those populations.

## METHODS

### Simulations model overview

All simulations were conducted in SLiM version 4.0.1 (Haller and Messer 2017; Haller and Messer 2023) with modifications to simulate tetraploids as done in Booker and Schrider (2025), where the full details of our simulation modifications can be found. Briefly, because SLiM does not support polyploids natively, we created polyploids by initializing two populations where one population functions as a storage population for the extra two chromosomes in each tetraploid, and the other population stores the other two chromosomes and is used to track the demographic and selection processes throughout the simulation. All polyploids were modeled as tetrasomic autotetraploids, with chromosomes only forming bivalents during meiosis (i.e. chromosomes pair up for recombination and random segregation during meiosis, rather than in groups of 3 or 4). When forming bivalents, all possible pairings of the 4 chromosomes were equally likely. For these simulations there was no burn-in period as this period is generally intended to get populations to equilibrium conditions, and the purpose of this study is to describe the nature of reaching equilibrium and if that equilibrium condition is pseudo-overdominance.

### Demographic and fitness model and associated parameters

Simulations were conducted under a constant-size Wright-Fisher (WF) model. Individuals in the simulation have a single chromosome 1Mb in length, and the per-base pair mutation and recombination rates are constant across all sites. For all simulations, population size *N* is constant and an individual’s fitness relative to that of all individuals in the population determines the probability of that individual being chosen for reproduction. Fitness is calculated multiplicatively according to Eq. 1:

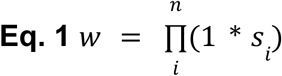

Where *w* is an individual’s fitness, and *s* is the selection coefficient for a given homozygous mutation *i.* For all simulations other than those using the empirical *h-s* DFE described below, mutations were fully recessive and *s* was constant across all mutations in a given simulation.

To incorporate more realistic parameters of dominance and selection coefficients, we also conducted simulations where mutations are drawn from an empirically derived DFE with a described relationship between dominance and selection coefficients (hereafter referred to as empirical *h-s* DFE). The DFE and *h-s* relationship was previously described from empirical data in *Arabidopsis lyrata* (Huber et al. 2018), and we used the equations from Booker and Schrider (2025) for translating discrete diploid dominance coefficients into continuous, ploidy independent coefficients. Eq. 2 describes the calculation of *h* from *s* given the value of *h* at s = 0 (*θ*_intercept_), and the rate at which *h* approaches 0 with a declining negative *s* (*θ*_rate_). The DFE for *s* was parameterized as a gamma distribution with *α* = 0.16 and *β* = 0.0092, multiplied by −1 to make mutations deleterious, and *h* was parameterized with *θ*_intercept_ = 0.978 and *θ*_rate_ = 50328. Importantly, because *s* in Eq. 2 is always negative, *h* is always between 0 and 1.

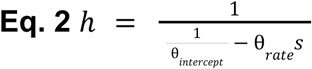

We varied all parameters of the simulation and analyzed multiple combinations of population size (*N*), selection coefficient (*s*), recombination rate (⍴), and the mutation rate (*μ*) to understand the interplay of these parameters in driving pseudo-overdominance in diploid and tetraploid populations (see Table 1). Distributions of these parameter values were chosen based on initial tests to capture the range of values where pseudo-overdominance could occur given the other fixed parameters. Due to the large number of parameter combinations, simulations were limited to 10 replicates per unique set of parameters.

**Table 1.** Parameters and their sampled values used for the simulation.

| Parameter | Values |
| --- | --- |
| population size ($N$) | 100,200,500 |
| selection coefficient ($s$) | -0.0005,-0.001,-0.005,-0.01,-0.02,<br>empirical $h$-s DFE |
| recombination rate ($\rho$) | 1e-10,1e-09,5e-09,1e-08,5e-08,1e-07 |
| mutation rate ($\mu$) | 1e-10,1e-09,5e-09,1e-08,5e-08,1e-07($N=100$) |
| ploidy | autopolyploid ($4n$), diploid ($2n$) |

We note the simulation model described above is not meant to approximate the dynamics of selection and pseudo-overdominance in realistic populations or genomes, but rather explore the parameter space in which pseudo-overdominance is more likely to occur in autotetraploids than diploids.

### Describing Pseudo-overdominant dynamics

To characterize pseudo-overdominant dynamics, we focus on four main statistics. As previously described, pseudo-overdominance results in intermediate median allele frequencies due to the maintenance of distinct haplotypes with non-complementary deleterious mutations (Gilbert et al. 2020; Zhao and Charlesworth 2016; Pamilo and Pálsson 1998). In addition, because pseudo-overdominance favors heterozygous individuals whose recessive deleterious alleles are masked, we would expect it to slow the reduction in the population’s mean fitness as deleterious mutations accumulate. Finally, because overdominance impedes the fixation of selected (and linked) alleles, these deleterious mutations should present as polymorphisms rather than fixations. These expectations are borne out by our results shown in Figs. 1–2: population fitness levels off or declines less rapidly in populations experiencing psuedo-overdominance, and deleterious segregating mutations continually accumulate in the form rather than being purged by selection, while the rate of fixation stagnates. Given these statistics, we then describe pseudo-overdominance as occurring when populations experience: (1) elevated intermediate median allele frequencies and (2) decelerated population fitness decline while also experiencing (3) a continual increase in the number of segregating mutations, but (4) do not experience a concordant increase in the number of fixed mutations at a similar rate. Generally, the combination of all of these is indicated by elevated intermediate median allele frequencies, but some scenarios where this is not the case are discussed.

**Figure 1.**
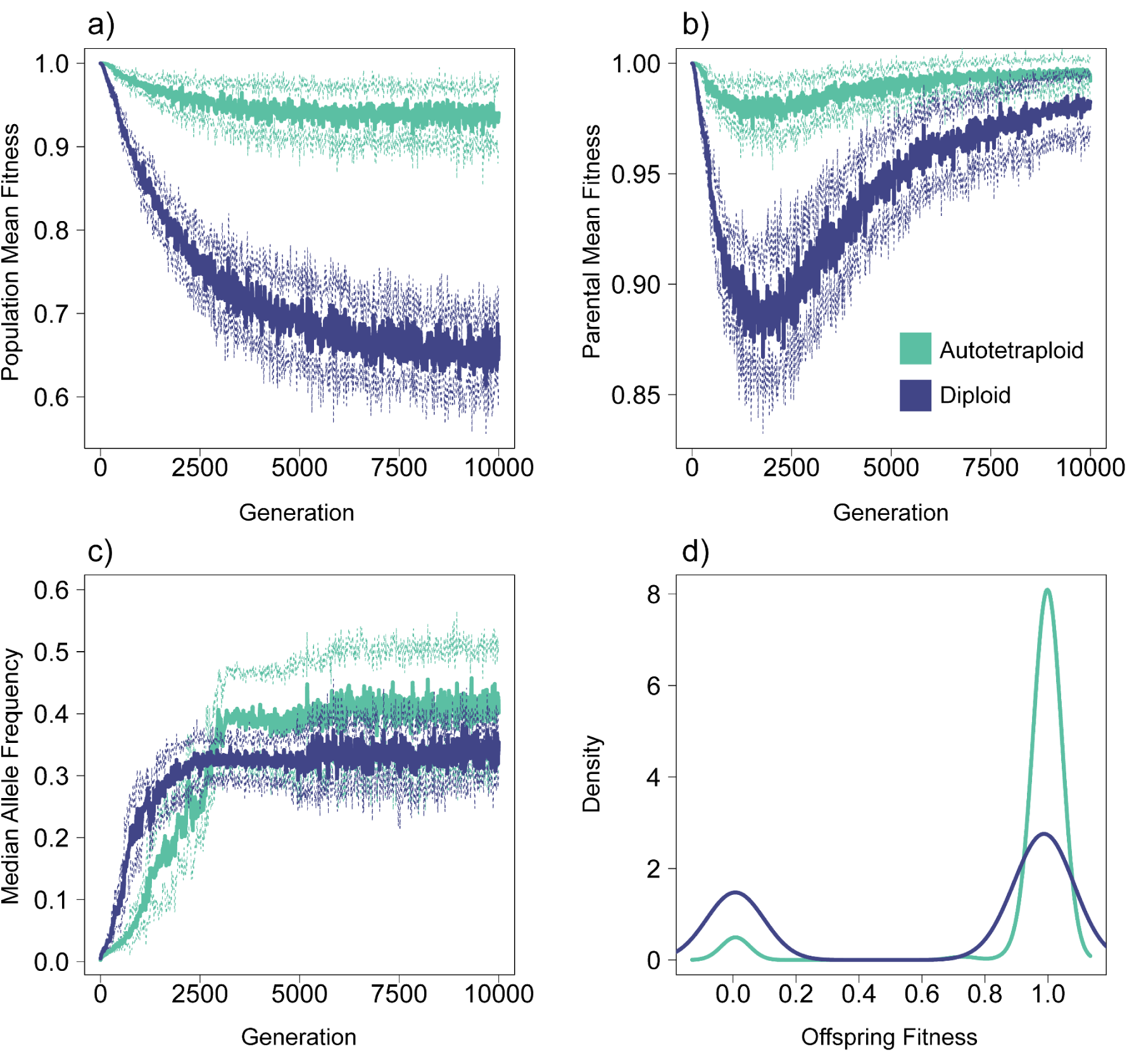
Differences between diploids and autotetraploids under pseudo-overdominant dynamics. a) Mean fitness of the total population at each generation. Solid lines represent the average across 10 simulations, with dashed lines representing ±1 standard deviation. Note that these values are not relative fitness, but SLiM’s absolute fitness calculation based on all segregating deleterious mutations present in an individual. b) Mean fitness of parents chosen for reproduction at each generation. c) Median allele frequency at each generation. d) Density of offspring fitness for diploids and tetraploids at generation 10,000 of the simulation. The parameters for the simulation replicates shown in this figure are *N* = 100, *μ* = 1.0e-07, *s* = −0.005, ⍴ = 1.0e-10.

**Figure 2.**
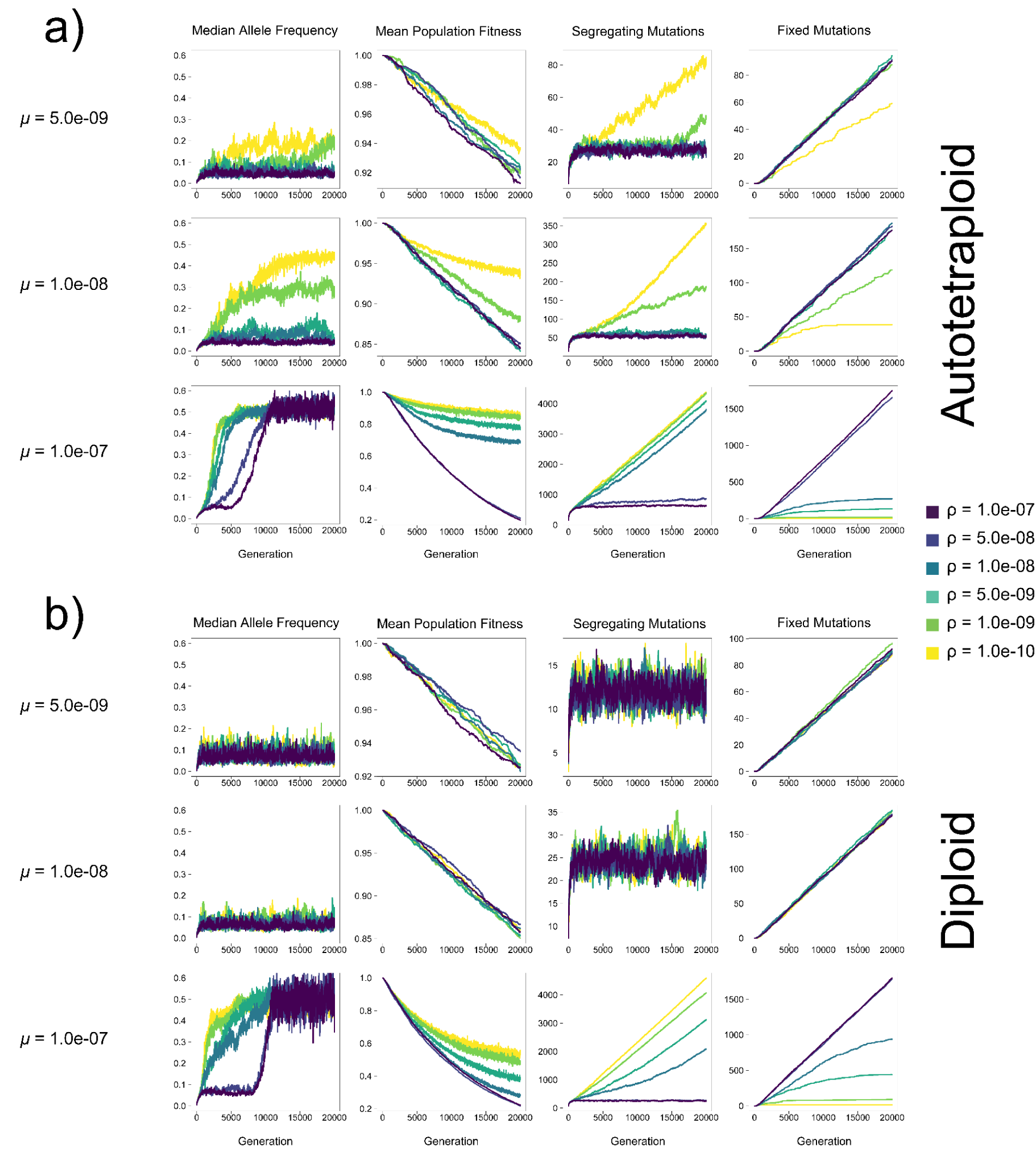
Median allele frequency, mean population fitness, the number of segregating mutations, and the number of fixed mutations in the population at each generation for autotetraploids (top) and diploids (bottom) in simulations under the empirical *h-s* DFE with *N* = 100 individuals. Each line represents the average across 10 replicates.

## RESULTS

### Fitness is stable and high in autotetraploids under pseudo-overdominant dynamics

We first characterize and describe the differences between autotetraploids and diploids under pseudo-overdominant dynamics. Using an illustrative example (*N* = 100, *μ* = 1.0e-07, *s* = −0.005, ⍴ = 1.0e-10), the fitness of both autotetraploids and diploids decreases prior to reaching equilibrium pseudo-overdominant dynamics (Fig. 1a); however, population fitness decreases much more in diploids and continues to decrease well after pseudo-overdominant dynamics are occurring (∼ generation 2000, inflection point in parental mean fitness curve, Fig. 1b). In contrast to the overall population fitness, Fig. 1b shows that the fitness of individuals sampled for reproduction recovers after average allele frequencies reach equilibrium dynamics (∼ generation 2500 here, Fig. 1c) in both diploids and autotetraploids, despite the fact that the fitness difference between heterozygous and homozygous individuals continues to widen after this point (as seen in the different fitness trajectories for the population as a whole and those selected to be parents in Fig. 1a,b). The cause of the difference in fitness between ploidies under pseudo-overdominance is evident from the distributions of offspring relative fitness values in autotetraploids and diploids (Fig. 1d). While eventually the accumulation of mutations results in fitness approaching zero in homozygotes of both diploids and tetraploids, the distribution of offspring genotypes in the two groups is drastically different, with a much smaller proportion of low-fitness homozygotes in tetraploids. The higher average allele frequency in tetraploids may partially account for this, but the smaller probability of forming homozygotes in tetraploids plays a greater role. For example, with an allele frequency of 0.4 (which is the approximate median at equilibrium for tetraploids in Fig. 1c) we would expect 52% of offspring genomes to be homozygotes in a diploid population vs. ∼16% in a tetraploid population.

### Pseudo-overdominant dynamics occur over a wider range of parameters in autotetraploids than in diploids

We performed simulations of diploid and autotetraploid populations under a variety of combinations of *μ*, ⍴, *s*, across three different population sizes (*N*=100, *N*=200, and *N*=500). We then asked whether these simulations met four criteria for pseudo-overdominance: 1) intermediate allele frequencies (examined in Fig. 4, Supp. Figs. 1-9), 2) an excess of deleterious polymorphisms (Fig. 5, Supp. Figs. 10-15), 3) an attenuation in the fitness decline as these deleterious polymorphisms accumulate (Fig. 6, Supp. Figs. 16-21), and 4) a diminished rate of fixation (Fig. 7, Supp. Figs. 22-27). The rationale for these criteria is shared in the Methods. Our simulation results show that in both autotetraploids and diploids pseudo-overdominance occurs more often when ⍴ is low, *μ* is high, and or |*s*| is high, but the relative area of parameter space where the four criteria of pseudo-overdominance are met is far larger in autotetraploids than in diploids (Figs. 2–7, Supp. Figs. 1-27). We note that there were some cases where intermediate median allele frequencies were observed but the other criteria were not met. For example, a small subset of high-recombination rate models showed elevated median allele frequencies but no concomitant accumulation of segregating mutations or decrease in the rate of fixation, along with a dramatic decrease in mean population fitness (e.g. the cases where *μ* = 1.0e-07 and ⍴ = 1.0e-07 in, Fig. 2). These parameter combinations therefore do not appear to be truly producing pseudo-overdominance. However, we did observe that for those parameter combinations where the median allele frequency was elevated in autotetraploids but not diploids, the other three criteria for pseudo-overdominance were typically met in autotetraploids. This suggests that a comparison of median allele frequencies (averaged across replicates) in autotetraploids vs. diploids serves as a useful proxy for identifying regions of the parameter space where pseudo-overdominance is more likely in the former. We show these in the form of heatmaps (Fig. 3 and Supplementary Figures 1-3) which can be used to summarize the relative propensity for pseudo-overdominance in the two ploidy levels, and which underscore the much larger region of parameter space where pseudo-overdominance emerges in autotetraploids. Moreover, these heatmaps may actually underestimate the extent to which pseudo-overdominance is more prevalent in autotetraploids, as there are some parameter combinations where autotetraploids meet all four of our criteria but diploids only exhibit an elevation in median allele frequencies (e.g. *N*=100, *μ*=5e-8, *s*=-0.001, and ⍴=1e-8, Supp. Figs. 4–5, 10-11, 16-17, 22-23).

**Figure 3.**
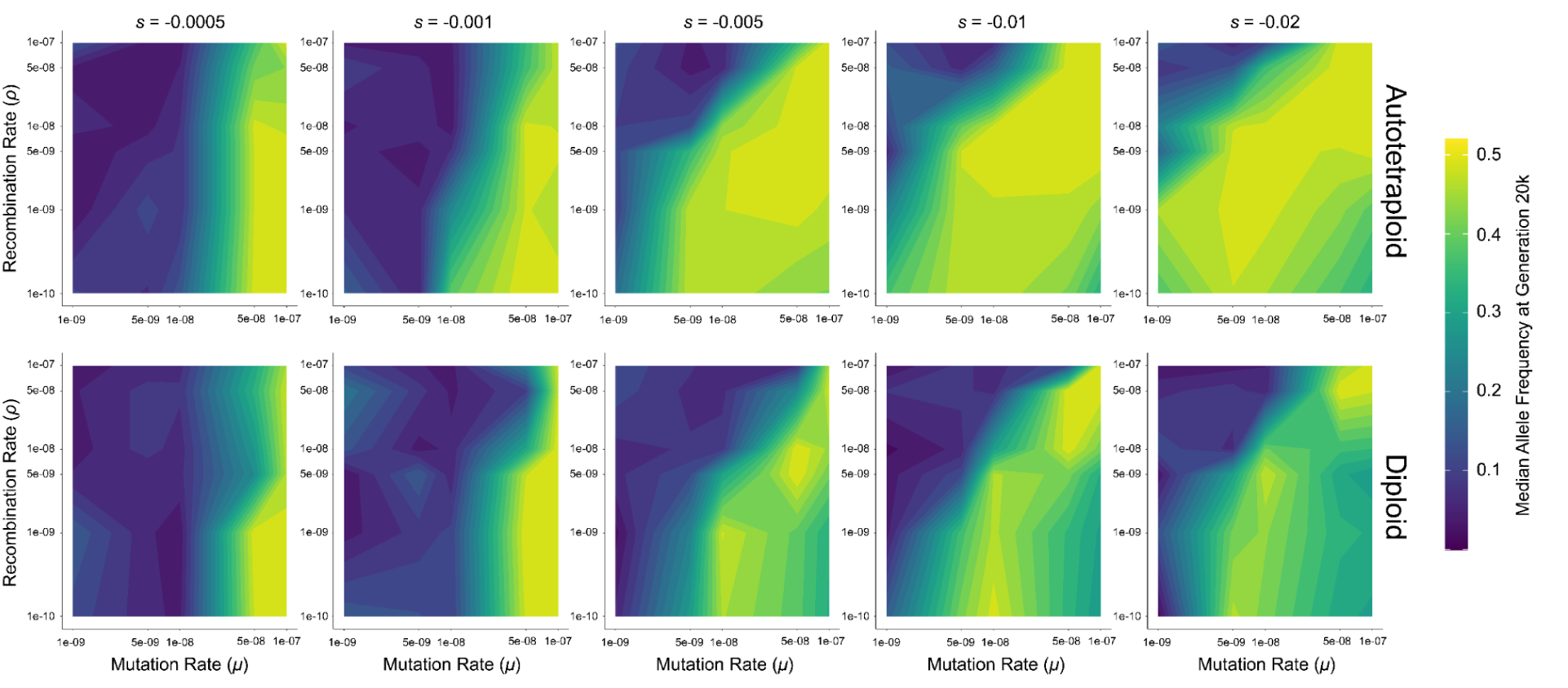
Median allele frequency at 20,000 generations for autotetraploid (top) and diploid (bottom) simulations of a population of *N* = 100 individuals. Axes are transformed to log scale, with values between ticks interpolated to fill in the gradient. Allele frequencies for each parameter combination are averaged across 10 replicates.

**Figure 4.**
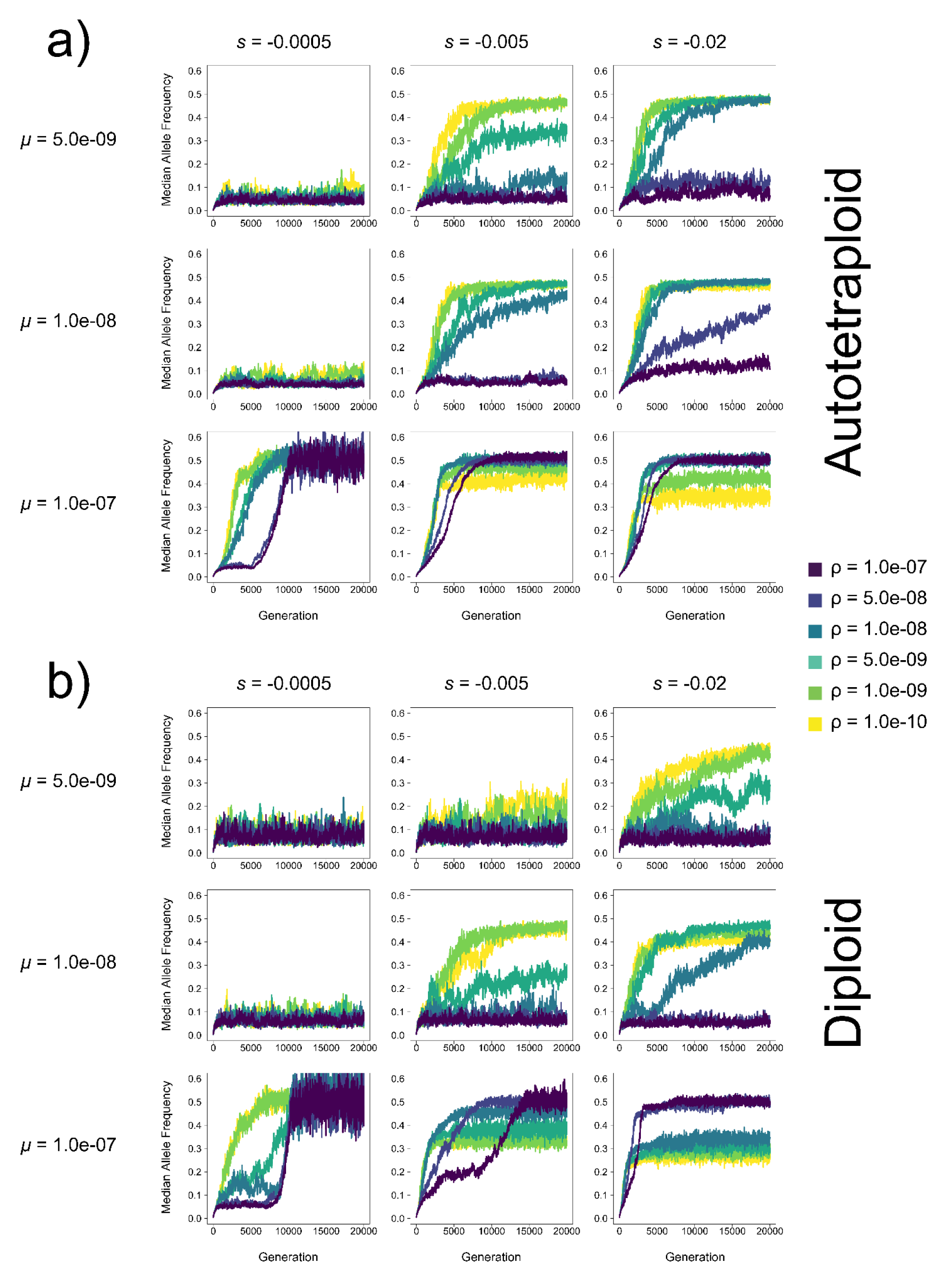
Median allele frequency of (a) autotetraploids and (b) diploids in each generation at varying mutation rates (*μ*) and selection coefficients (*s*) for a population of *N* = 100 individuals. Each line represents the average across 10 replicates.

**Figure 5.**
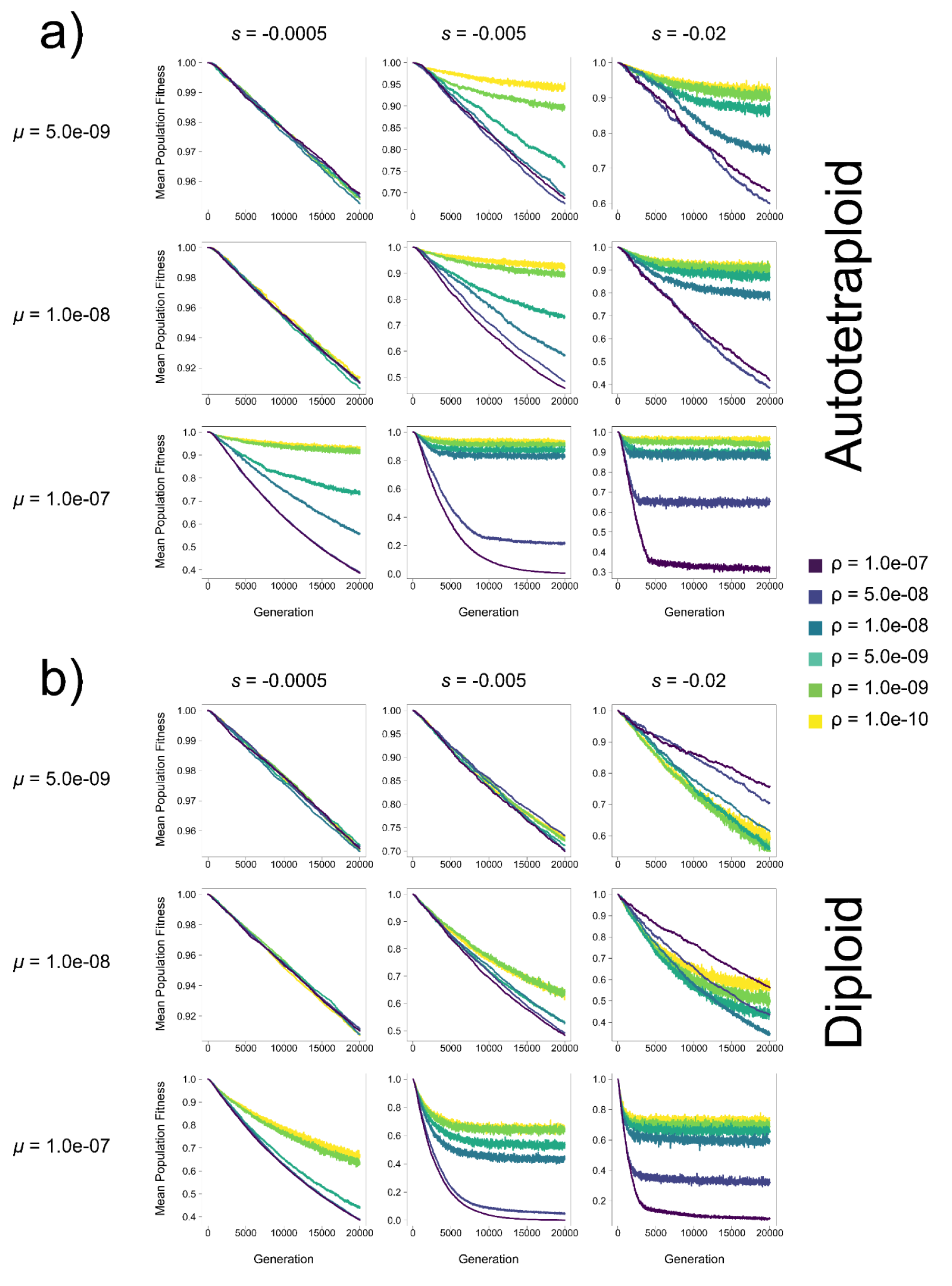
Mean population fitness of (a) autotetraploids and (b) diploids in each generation at varying mutation rates (*μ*) and selection coefficients (*s*) in a population of *N* = 100 individuals. Each line represents the average across 10 replicates. Note: y-axis varies across subfigures.

**Figure 6.**
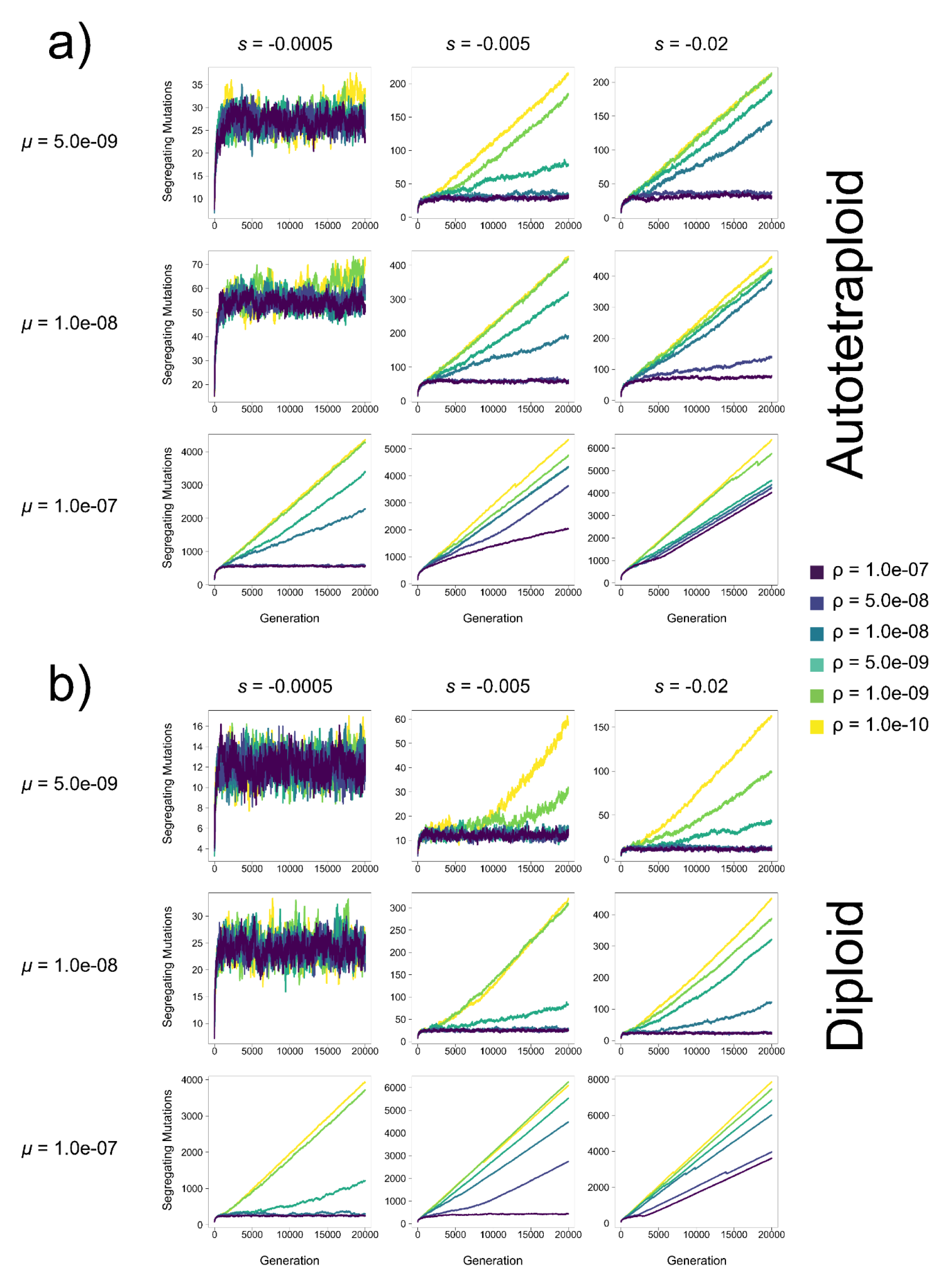
The number of segregating mutations in populations of (a) autotetraploids and (b) diploids in each generation at varying mutation rates (*μ*) and selection coefficients (*s*) in a population of *N* = 100 individuals. Each line represents the average across 10 replicates. Note: y-axis varies across subfigures.

**Figure 7.**
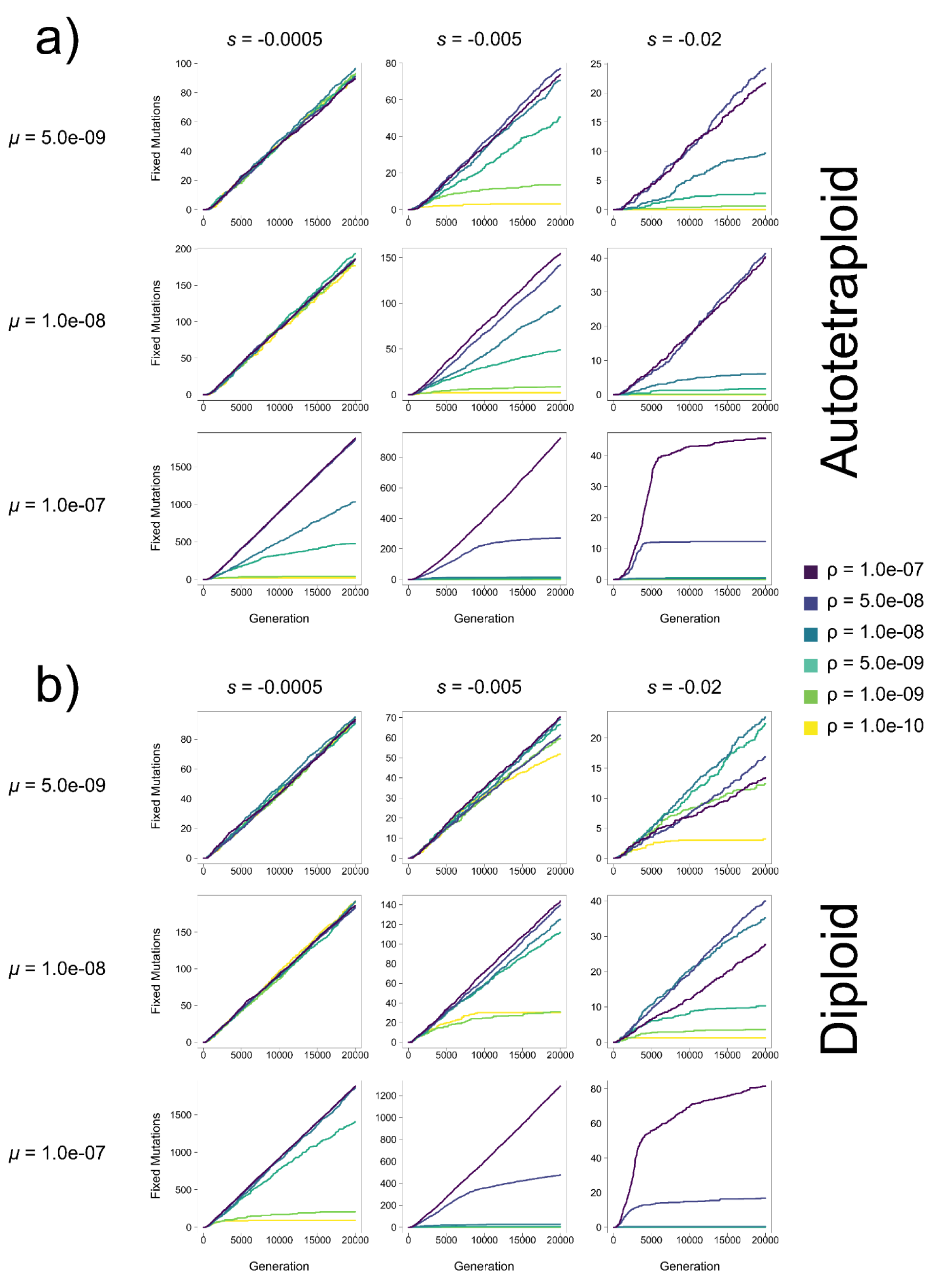
The number of fixed mutations in populations of (a) autotetraploids and (b) diploids in each generation at varying mutation rates (*μ*) and selection coefficients (*s*) in a population of *N* = 100 individuals. Each line represents the average across 10 replicates. Note: y-axis varies across subfigures.

The above results were obtained from relatively simplified models where all deleterious mutations are fully recessive. To determine whether this holds for more realistic distributions of dominance coefficients, we repeated our simulations under a distribution of deleterious fitness effects estimated from empirical data by Huber et al. (2018). Importantly, this DFE contains variation in dominance coefficients, with *h* varying according to the value of *s*. We observed qualitatively similar results under this model as under our *h*=0.0 simulations above: pseudo-overdominance is more likely when the recombination rate is low, and/or deleterious mutation rate is high, and is more common in autotetraploids than diploids (Fig. 2 and Supplementary Figure 1).

Additionally, although in most cases the repulsion effects of pseudo-overdominance when present are obvious across all criteria simultaneously and early on in our simulations, in some parts of parameter space the consequences of pseudo-overdominance and its signatures are more subtle. For instance, in autotetraploids where *μ* = 1.0e-07, *s* = −0.0005, and ⍴ = 5.0e-09, the number of fixed mutations does initially increase, as is the case for simulations without pseudo-overdominance. However, this increase is slower than that of populations with higher recombination rates, and appears to be tailing off prior to the end of the simulation (Fig. 7), while similarly the number of segregating mutations is initially slow but accelerates during the simulation (Fig. 6). This is mirrored by an initial decline in mean population fitness, followed by a delayed but noticeable deceleration of this decline (Fig. 5). While this population may not be in total pseudo-overdominant repulsion here by the end of the simulation, the evolutionary dynamics that eventually result in that repulsion are clearly acting upon that population. This is particularly evident when looking at median allele frequencies which increase rapidly and early (Fig. 4), suggesting an evolutionary drive to maintain alternative alleles in the population even if upon termination of the simulation some the population is still at intermediate (but rising) values for some of our criteria for pseudo-overdominance.

Finally, we found that for some parameter combinations, pseudo-overdominance appeared to be more likely in larger populations than smaller ones. While this was observed in both autotetraploids and diploids, we highlight only diploid examples to emphasize that this phenomenon is not limited to higher ploidies. At lower values of *s* and ⍴, evidence of pseudo-overdominance appears at low mutation rates when *N* = 500 but not when *N* = 100 (e.g. *μ* = 5.0e-09, *s*=-0.0005, and ⍴ = 1.0e-10 in diploids at N=500; Fig. 3, Supp. Figs. 2-3,5,9,11,15,17,21,23,27). In some regions of the parameter space with higher mutation rates (e.g. *μ* = 5.0e-08, *s*=-0.001, and ⍴ = 5.0e-09 in diploids) there are subtle trends toward pseudo-overdominance at *N* = 100 (Supp. Figs. 5,11,17,23) but stronger evidence that our four criteria are being met at *N* = 500 (Supp. Figs. 9,15,21,27). We observe similar trends when comparing *N* = 200 to *N* = 500 when *μ*=1e-08, *s*=-0.0005, and ⍴ = 1.0e-10. While it is possible that the smaller-*N* populations in the examples above will eventually reach repulsion, *N* = 500 populations in this space do so rapidly for each of our four criteria for pseudo-overdominance. Together these results indicate that in some parts of our parameter space pseudo-overdominance indeed occurs more readily in large populations, contrary to prior expectations.

### Lower recombination rates result in higher fitness under pseudo-overdominance

A striking result demonstrating the fitness benefits underlying pseudo-overdominance is that for several combinations of *μ*, *s*, and *N* the recombination rate is inversely proportional to fitness in both diploids and autotetraploids, and the range of parameter space where this fitness-recombination relationship occurs is much greater in autotetraploids (Fig. 5, Supp. Figs. 10-15). Furthermore, for autopolyploids in any specific combination of *N*, *s*, and *μ*, where pseudo-overdominance does occur at some ⍴, a reduction in ⍴ is either selectively neutral or advantageous in nearly all cases. The same is not true for diploids, where there is often a fitness cost in reducing ⍴ from 1.0e-7 (1 recombination per chromosome per generation) to an intermediate ⍴ before the benefits of pseudo-overdominance are realized in lower recombination rates (e.g. high *s* simulations in Supp. Figs. 11,13,15); while similar trends are seen in some instances for autotetraploids, the differences in fitness across recombination rates are much smaller in magnitude in such cases, and are frequently only realized after several thousands of generations (e.g. *μ*=5e-9 and *s*=-0.02 in Supp. Figs. 14, 15). In total, this result illuminates the routes to pseudo-overdominance in both diploids and autotetraploids but demonstrates that the path is clearer in autotetraploids.

## Discussion

### Pseudo-overdominance is expected to be far more prevalent in polyploids than diploids

As evolutionary processes, pseudo- and associative overdominance are generally considered to be rare. This is at least in part because in order for diploids populations to enter pseudo-ovderominant dynamics the values of several population genetic parameters must each be within a relatively narrow range (i.e. sufficiently low recombination, recessive mutations, and large enough cumulative deleterious fitness effects). Thus, there is an especially small region of the joint parameter space in which all of these conditions can be satisfied (Charlesworth and Jensen 2021). Gilbert et al. (2020) showed that under a multi-locus model, this parameter space is wider than previously thought. Here we show that for organisms of higher ploidy the range of conditions where pseudo-overdominance is expected is dramatically expanded. As such, pseudo-overdominance may be a much more prominent evolutionary force than currently appreciated.

The main cause for the increased relevance of pseudo-overdominance in polyploids is rather straightforward: the masking effect of additional allelic copies allows polyploid organisms to tolerate a greater load of recessive deleterious mutations—effectively lowering the barrier of entry into pseudo-overdominant parameter space. As demonstrated here, fitness remains high in polyploids both prior to and after entering into pseudo-overdominant dynamics (Fig. 1). Conversely, in diploids the transition to pseudo-overdominant equilibrium is preceded by a dramatic drop in fitness, and after the transition even those individuals selected to reproduce exhibit much lower fitness than observed in pseudo-overdominant polyploid populations.

### The interplay between pseudo-overdominance and recombination rate

Results from our simulations demonstrate that for both diploids and polyploids, there are combinations of *μ*, *s*, and *N* where population fitness appears to be inversely proportional to the recombination rate (Fig. 5, Supp. Figs. 10-15). Furthermore, this was observed in across a much wider range of the parameter space in polyploids, with the majority of *μ*, *s*, and *N* combinations that elicited pseudo-overdominance also demonstrating this inverse relationship. This is consistent with theoretical results demonstrating that pseudo-overdominance occurs when the relative absence of recombination prevents high-load alleles from exchanging deleterious mutations across haplotypes resulting in higher fitness heterozygotes despite that greater load (Frydenberg 1963; Ohta and Kimura 1970; Ohta 1971; Charlesworth 1991).

While our results are largely consistent with the notion that pseudo-overdominance should be more prevalent in smaller populations, this does not always appear to be the case. We observe that when selection coefficients and recombination rates are low (e.g. toward the bottom of the plots shown leftmost columns of Fig. 3 and Supp. Figs. 2-3) pseudo-overdominance emerges more readily when *N*=200 or *N*=500 than when *N*=100; this is the case for both diploids and autotetraploids. The explanation for this is simply that, when *s* is very low, a larger number of deleterious mutations are needed to produce haplotypes with large enough fitness effects such that pseudo-overdominance may arise. This could be achieved by increasing *μ*, or instead by increasing the population size, thereby increasing the population-scaled mutation rate parameter *θ*=4*Nμ*. This will not result in pseudo-overdominance when *r* is moderate or high, because increasing *N* also increases the population-scaled recombination rate *ρ*=4*Nr*, thereby preventing the build-up of haplotypes with complementary sets of deleterious alleles. Thus, only when *r* is very low might we expect a larger population to have a greater propensity for pseudo-overdominance resulting from the accumulation of weakly deleterious alleles.

Previous work on pseudo-overdominance has largely focused on the genomic conditions that support the buildup of deleterious mutations and linked neutral diversity to enter a pseudo-overdominant state, and evidence of pseudo-overdominance has been found or been suggested to occur in some organisms with genomic regions reflecting these conditions (Becher et al. 2020; Gilbert et al. 2020). However, this relationship may be bidirectional, as one might expect pseudo-overdominance to impact the forces that shape recombination rates as well. Indeed, it has been shown that mutations that increase the local recombination rate would be disfavored under pseudo-overdominance (Pálsson 2002), potentially implying that recombination suppressors could be favored. While we did not directly investigate whether recombination modifying alleles would successfully invade under pseudo-overdominant conditions, our results do suggest a clear path through which lower recombination could evolve due to pseudo-overdominance without having to pass through any intermediate recombination rate fitness valley. Moreover, our results suggest that any relationship between pseudo-overdominance and selection on recombination modifiers would be particularly pronounced in polyploids, and that this relationship may contribute to the consistently observed recombination suppression observed in newly formed polyploids (Morrison and Rajhathy 1960; Reddi 1970; Wu et al. 2014; Bomblies et al. 2016; Morgan et al. 2021).

### Variation in the timing and abruptness of the transition to pseudo-overdominant equilibrium

Pseudo-overdominance can only emerge when multiple complementary haplotypes each with a combined large deleterious fitness effect can form while evading selective purging of their constituent deleterious mutations. In many of the simulated scenarios we examined, the transition to pseudo-overdominant equilibrium occurs relatively rapidly once the simulation begins, suggesting that pseudo-overdominance may drive population dynamics immediately following the strong bottleneck of polyploid formation in genomic regions favoring these dynamics (e.g. see most of the low-recombination rate results in Fig. 4a). However, the transition can also occur gradually when the recombination rate is intermediate and intermediate combinations of *s* and *μ* (i.e. intermediate in both, or high *s* and low *μ*, or vice versa, exemplified in many of the diagonal or near-diagonal plots moving from bottom-left to top-right in Supp. Figs. 4,6). In such cases blocks of mutations may be sufficiently deleterious to be immediately subject to negative selection, but the recombination rate is not high enough for selection to remove these mutations as fast as they accumulate—an effect compounded by the masking effect of polyploidy.

### Evidence of pseudo-overdominance in polyploid species and crops

As discussed above, there are numerous conditions specific to polyploid formation that favor entering pseudo-overdominance: reductions in rho (Morrison and Rajhathy 1960; Bomblies et al. 2016; Morgan et al. 2021), and low census or effective population sizes (Stebbins 1950), perhaps through self-fertilization or periodic asexuality (Pálsson 2001; Barringer 2007; Stenberg and Saura 2013; Neiman et al. 2014). To date, pseudo-overdominance or associative overdominance has not been directly implicated in polyploids. However, there is some circumstantial evidence in salmonids that warrant further investigation: while investigating allele surfing, Rougemont et al. (2023) found that residual non-diploidized autotetraploid regions of the Coho salmon (*Oncorhynchus kitsuch*) genome harbored a far greater deleterious load than diploidized regions, and that this tetraploid-specific load was concentrated in genomic segments with low recombination rates. Furthermore, Bierne et al. (2000) suggested the correlation between heterozygosity and developmental stability in other *Oncorhynchus* trout (Leary et al. 1984) was evidence of associative overdominance in these species. Given these trout experienced the same tetraploidization event as Coho salmon and likely had a similar delayed diploidization, an investigation into the relationship between inheritance, genetic load, and the potential role of pseudo-overdominance in salmonids may be fruitful.

Although *Oncorhyncus* species have considerable economic value as game and food species, the potential role of pseudo-overdominance likely has more widespread agronomic implications, as numerous important crop species are polyploid (Renny-Byfield and Wendel 2014). For many such species, polyploidy itself was critical to the domestication process leading to dramatic phenotypic changes (Salman-Minkov et al. 2016; K. Zhang et al. 2019). However, the masking effect of polyploidy on recessive alleles also makes selection difficult as deleterious recessives are hard to remove from the population. Pseudo-overdominance may then compound this problem as any genomic regions under these dynamics will have an even higher genetic load, and high-load individuals can be highly fit but produce offspring with very low fitness. Furthermore, selection in crop breeding typically involves processes that work to reduce *N_e_* or promote heterosis and therefore breeders may inadvertently push more genomic regions into pseudo-overdominant dynamics thereby increasing the genetic load (for a discussion on heterosis and pseudo-overdominance, see Waller 2021).

Again, while pseudo-overdominance has not been directly attributed to genomic signatures in polyploids presently, circumstantial evidence in autopolyploid crops suggest its presence. Perhaps most clearly, genomic signatures and complications with selective breeding in potatoes are consistent with expectations from pseudo-overdominance. Compared with other crops, potatoes have had minimal gains in yield (Douches et al. 1996)—a phenomenon that may be attributable to difficulties in developing homozygous inbred lines in potato because of its high deleterious load (Lindhout et al. 2011). Although the production of self-compatible diploids has resulted in improvements, the residual load from the autotetraploids continues to plague breeders (Wu et al. 2023). Indeed, one inbred potato line retained highly heterozygous regions that accounted for 20% of the genome despite 9 generations of selfing (van Lieshout et al. 2020), and much less dramatic deviations from expected loss in heterozygosity from selfing in maize (which is diploid) were attributed to pervasive associative overdominance (Roessler et al. 2019). Zhang et al.’s (C. Zhang et al. 2019) investigation into the origin of inbreeding load in potatoes suggest most of the deleterious load is heterozygous and found in low recombination pericentromeric regions of the genome. Interestingly, much like our simulations showed pseudo-overdominance can occur in high recombination regions when mutations are sufficiently deleterious, Zhang et al. (C. Zhang et al. 2019) also found that the most strongly deleterious mutations (including lethals) occurred in high recombination regions and were found in a heterozygous state. Finally, Wu et al. (Wu et al. 2023) found that tetraploid potato lines harbored more heterozygous load than diploids, and their work suggest breeders should make selections based on maximizing homozygous load to prevent propagation of the more pervasive but masked heterozygous load.

Though the evidence is less clear than in potatoes, sugarcane is another crop where pseudo-overdominance may be responsible for difficulties in genetic gain. Modern sugarcane is a hybrid of octoploid *Saccharum spontaneum* and decaploid *Saccharum officinarum*, resulting in cultivars that vary in ploidy with between 8 and 12 chromosomal copies and recurrent aneuploidy (Amalraj and Balasundaram 2006). This initial hybridization of multiple species could have promoted pseudo-overdominance in addition to the elevated ploidy (Waller 2021), and although the present work focused on autotetraploids, the contribution of masking towards pseudo-overdominance due to additional chromosomal copies should be more pronounced in higher-level polyploids. Given the complex genome of sugarcane, uncovering strong population genetic evidence of pseudo- and associative overdominance would be challenging, but there are some indications of pseudo-overdominance in this highly polyploid species: Raboin et al. (2008) found pervasive high linkage disequilibrium across the sugarcane genome, and Ferreira et al. (2005) argued against selfing in sugarcane due to its high genetic load resulting in yield losses. Furthermore, the *S. spontaneum* genome has persisted despite repeated backcrossing with *S. officinarum* including the unreduced transmission of 2n *S. officinarum* chromosomes (Healey et al. 2024). Because yield gains have plateaued in sugarcane (Yadav et al. 2020), investigations into the role of pseudo-overdominance may provide insights that can help mitigate these deficiencies.

### Conclusions

Pseudo-overdominance occurs when selection is inefficient in purging recessive deleterious mutations due to small effective population sizes and genomic conditions that increase the disconnect between realized fitness and deleterious load. An increase in chromosomal copy number through whole genome duplication is one process which increases this disconnect. Although pseudo-overdominance is thought to be rare, multi-locus models have shown these dynamics can occur over a wider parameter space than previously appreciated, and our results show that this space is greatly expanded in polyploids. While we have highlighted a few examples of species with polyploid evolutionary histories that have demonstrated evidence of pseudo-overdominance, there are surely many more such cases in present-day polyploids, and recently diploidized genomes could also harbor signatures of past pseudo-overdominance. For example, divergence between duplicates may be elevated in regions previously evolving under pseudo-overdominant dynamics. Furthermore, pseudo-overdominance could drive the process of diploidization itself, resulting in older divergence times as the accumulation of complementary deleterious load favors the suppression of recombination and the divergence of haplotypes into distinct subgenomes. We have focused on crop species here due to the available evidence, but the implications for conservation—where small populations and hidden deleterious load are key concerns—also warrant consideration (Kyriazis et al. 2021). Similarly, although we have focused on autotetraploids as shorthand for polyploids with tetrasomic inheritance, our work is relevant for any polyploid with deviations from strict disomic inheritance—a common occurrence across both auto- and allopolyploids (Li et al. 2021). Ultimately, our work provides a basis for understanding pseudo-overdominant dynamics in polyploids, and more generally implies that pseudo-overdominance may be a more consequential evolutionary force than previously thought.

## Acknowledgements

We thank members of the Johri Lab for feedback on data generated for this project, and thank Austin Daigle for comments on the manuscript. WWB was supported by NIH award R01HG010774 and DRS was supported by NIH award R35GM138286.

## Data Availability

All simulation scripts can be found in the github repository https://github.com/wbooker/polyploid_pod.

## SUPPLEMENTAL INFORMATION

**Supplemental Figure 1.**
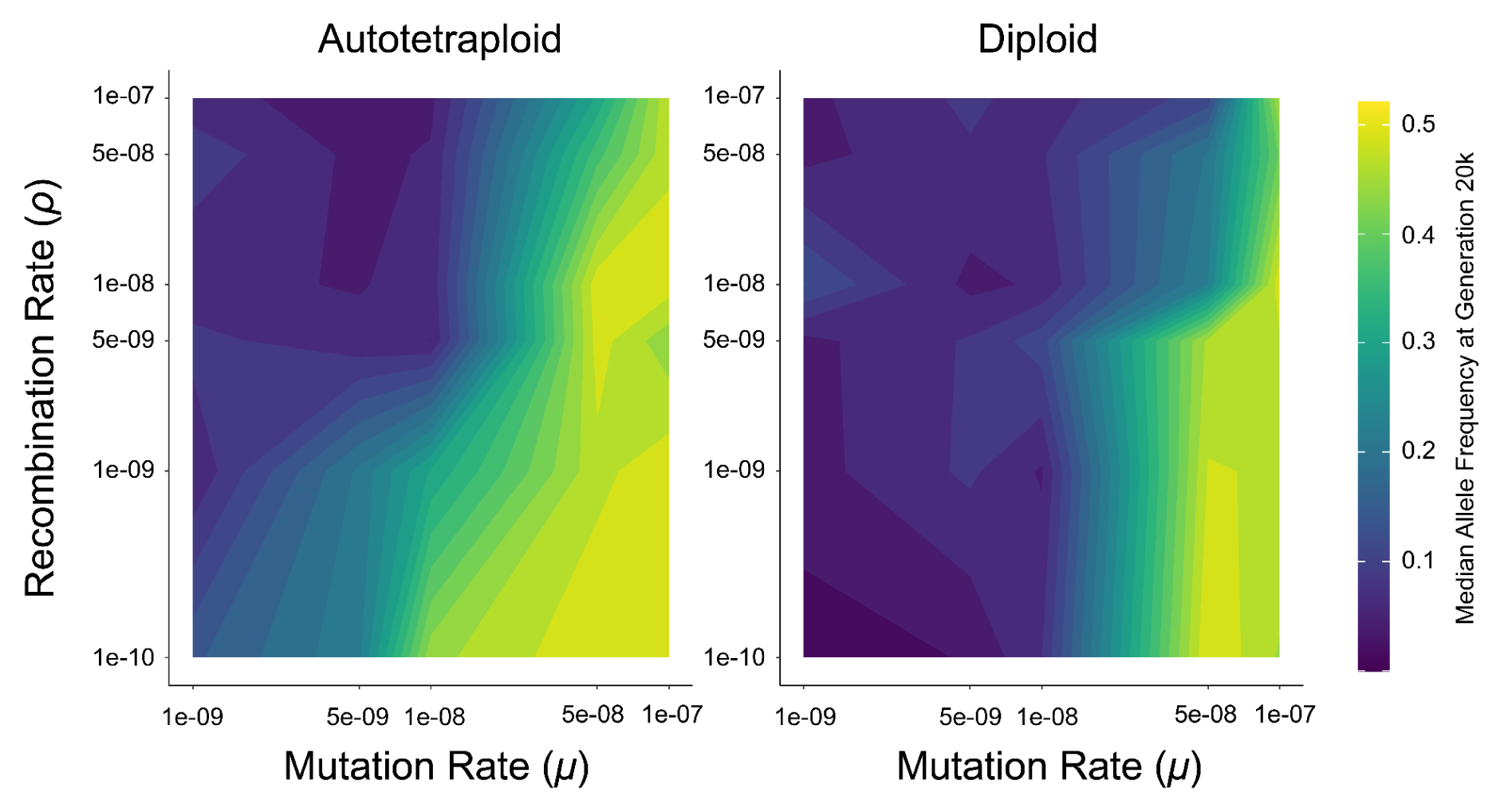
Median allele frequency at 20,000 generations for autotetraploid (top) and diploid (bottom) simulations under the empirical *h-s* DFE model with a population of *N* = 100 individuals. Axes are transformed to log scale, with values between ticks interpolated to fill in the gradient. Allele frequencies for each parameter combination are averaged across 10 replicates.

**Supplemental Figure 2.**
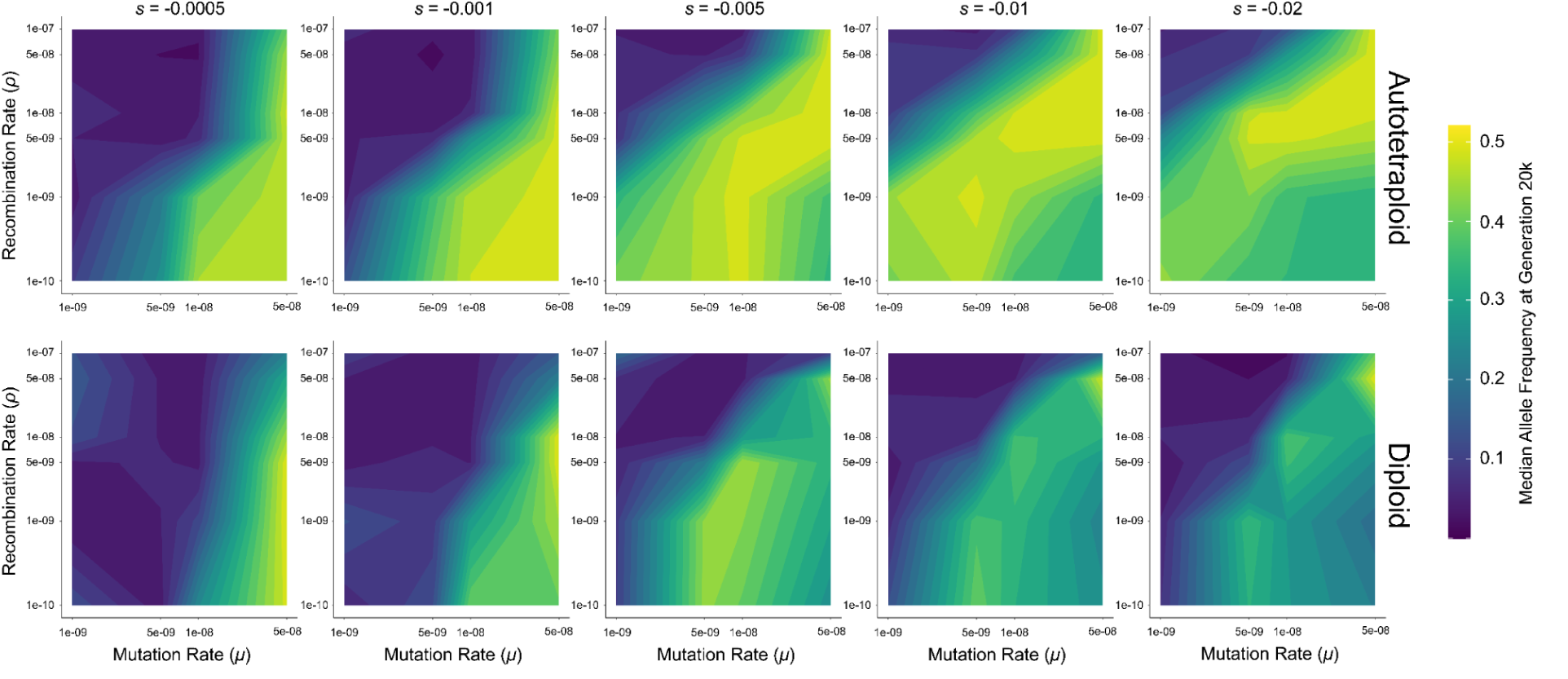
Median allele frequency at 20,000 generations for autotetraploid (top) and diploid (bottom) simulations of a population of *N* = 200 individuals. Axes are transformed to log scale, with values between ticks interpolated to fill in the gradient. Allele frequencies for each parameter combination are averaged across 10 replicates. Note, mutation rate at 1e-07 missing at this *N* as simulations were unable to finish due to the high mutation rate accumulating many mutations at low recombination rates.

**Supplemental Figure 3.**
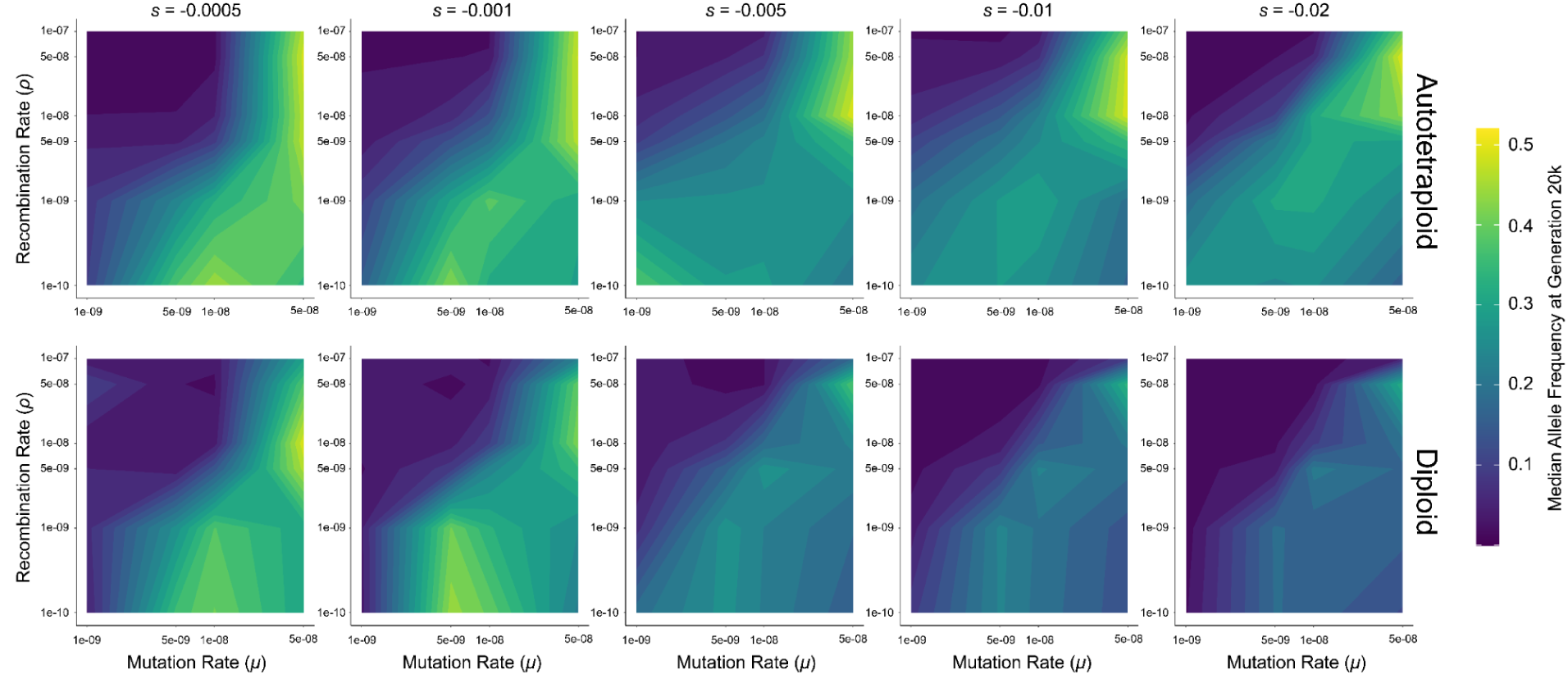
Median allele frequency at 20,000 generations for autotetraploid (top) and diploid (bottom) simulations of a population of *N* = 500 individuals. Axes are transformed to log scale, with values between ticks interpolated to fill in the gradient. Allele frequencies for each parameter combination are averaged across 10 replicates. Note, mutation rate at 1e-07 missing at this *N* as simulations were unable to finish due to the high mutation rate accumulating many mutations at low recombination rates.

**Supplemental Figure 4.**
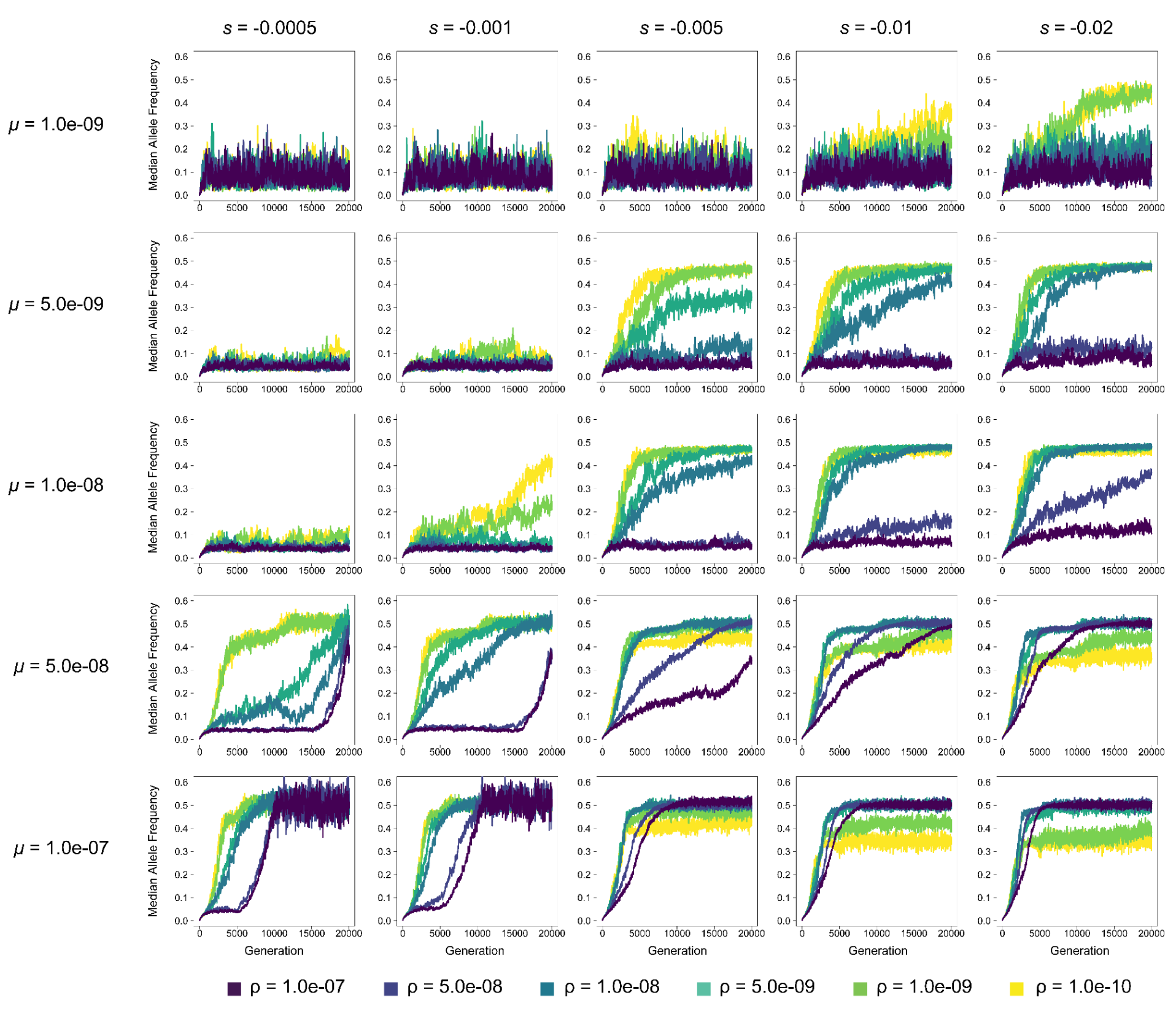
Median allele frequency of autotetraploids in each generation at varying mutation rates (*μ*) and selection coefficients (*s*) for *N* = 100. Each line represents the average across 10 replicates. Note: y-axis is identical across subfigures.

**Supplemental Figure 5.**
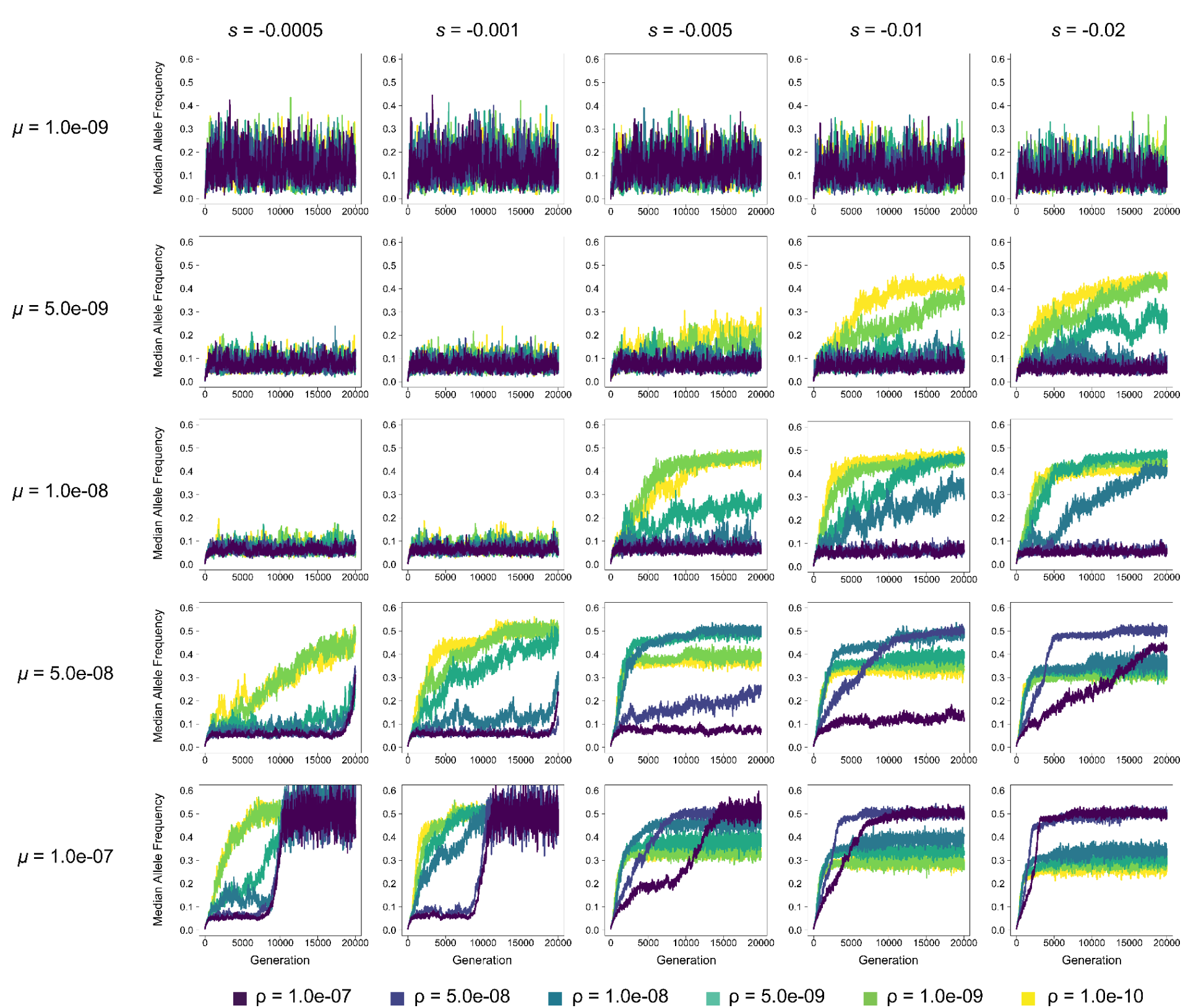
Median allele frequency of diploids in each generation at varying mutation rates (*μ*) and selection coefficients (*s*) for *N* = 100. Each line represents the average across 10 replicates. Note: y-axis is identical across subfigures.

**Supplemental Figure 6.**
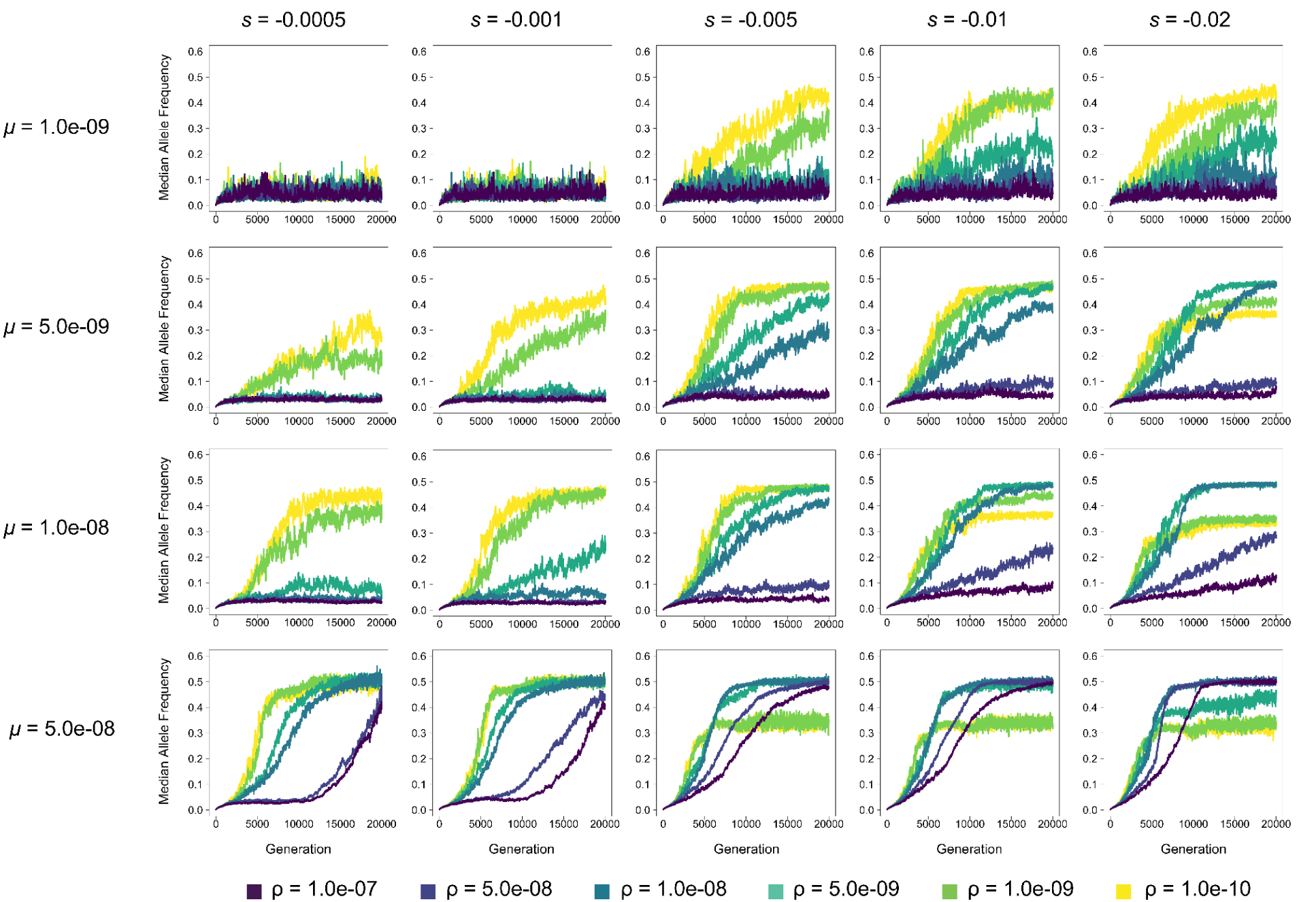
Median allele frequency of autotetraploids in each generation at varying mutation rates (*μ*) and selection coefficients (*s*) for *N* = 200. Each line represents the average across 10 replicates. Note: y-axis is identical across subfigures, and that *μ*=1e-7 simulations are not included here because these replicates could not complete in the maximum allotted time on our compute cluster for all *N*.

**Supplemental Figure 7.**
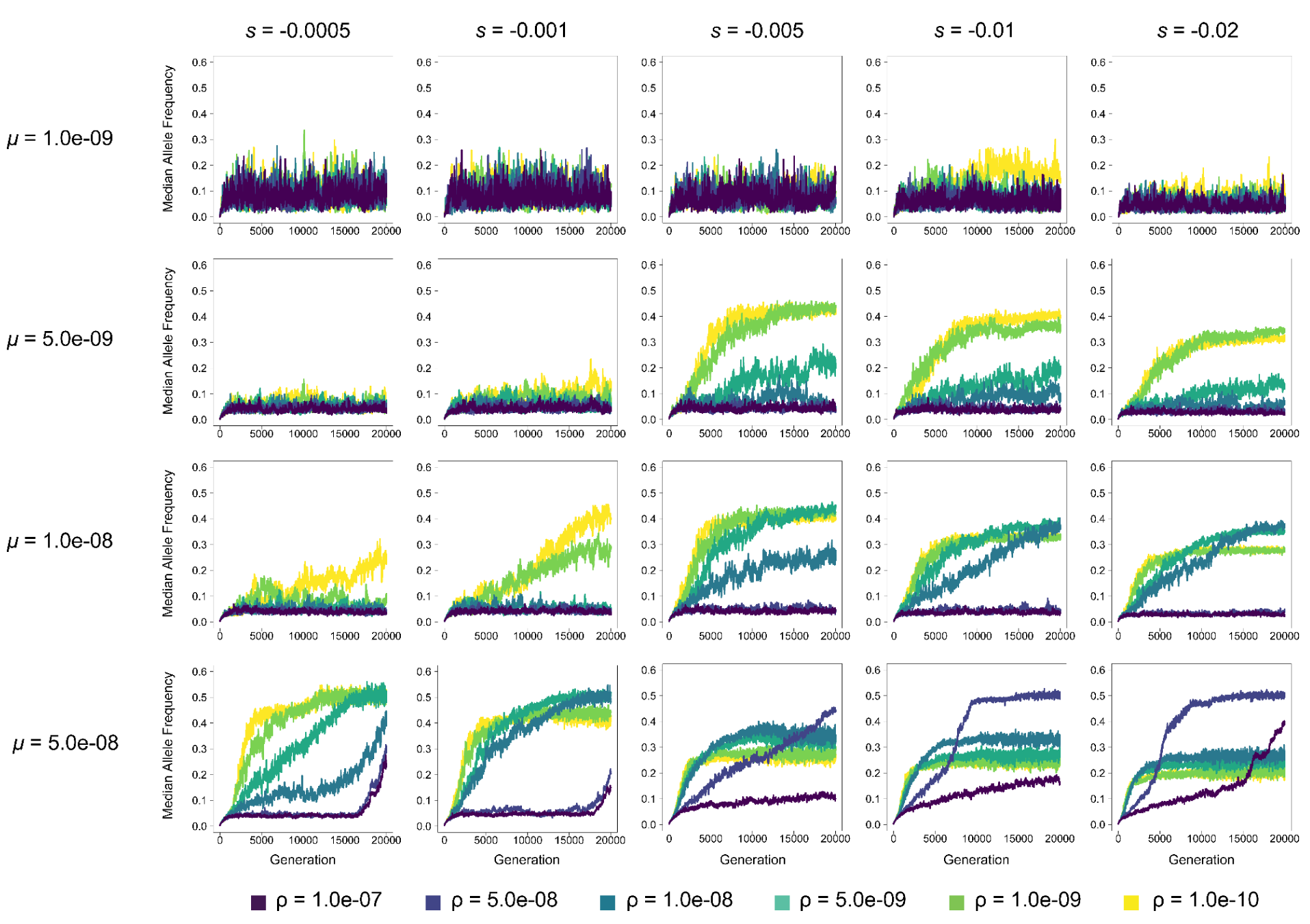
Median allele frequency of diploids in each generation at varying mutation rates (*μ*) and selection coefficients (*s*) for *N* = 200. Each line represents the average across 10 replicates. Note: y-axis is identical across subfigures.

**Supplemental Figure 8.**
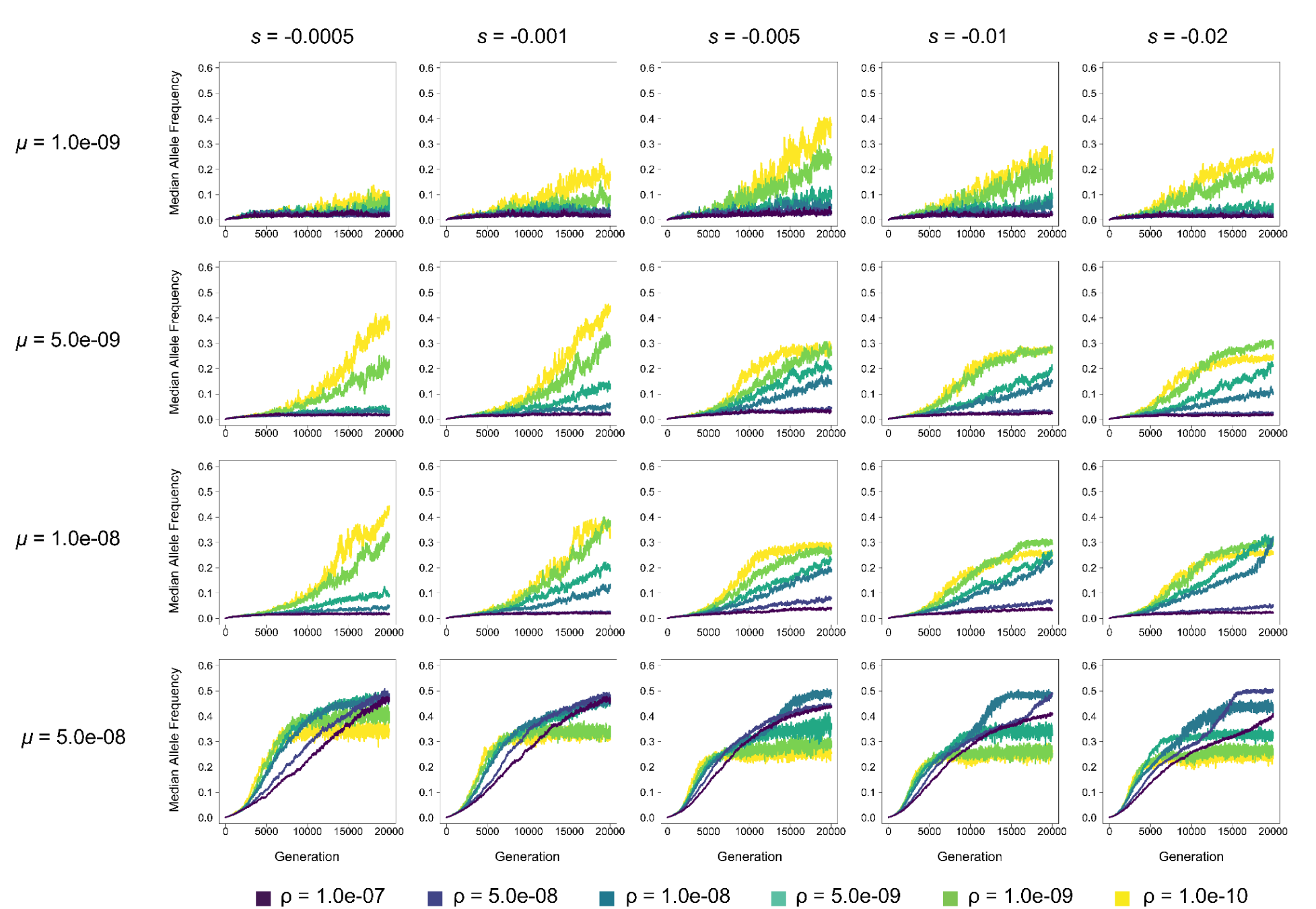
Median allele frequency of autotetraploids in each generation at varying mutation rates (*μ*) and selection coefficients (*s*) for *N* = 500. Each line represents the average across 10 replicates. Note: y-axis is identical across subfigures.

**Supplemental Figure 9.**
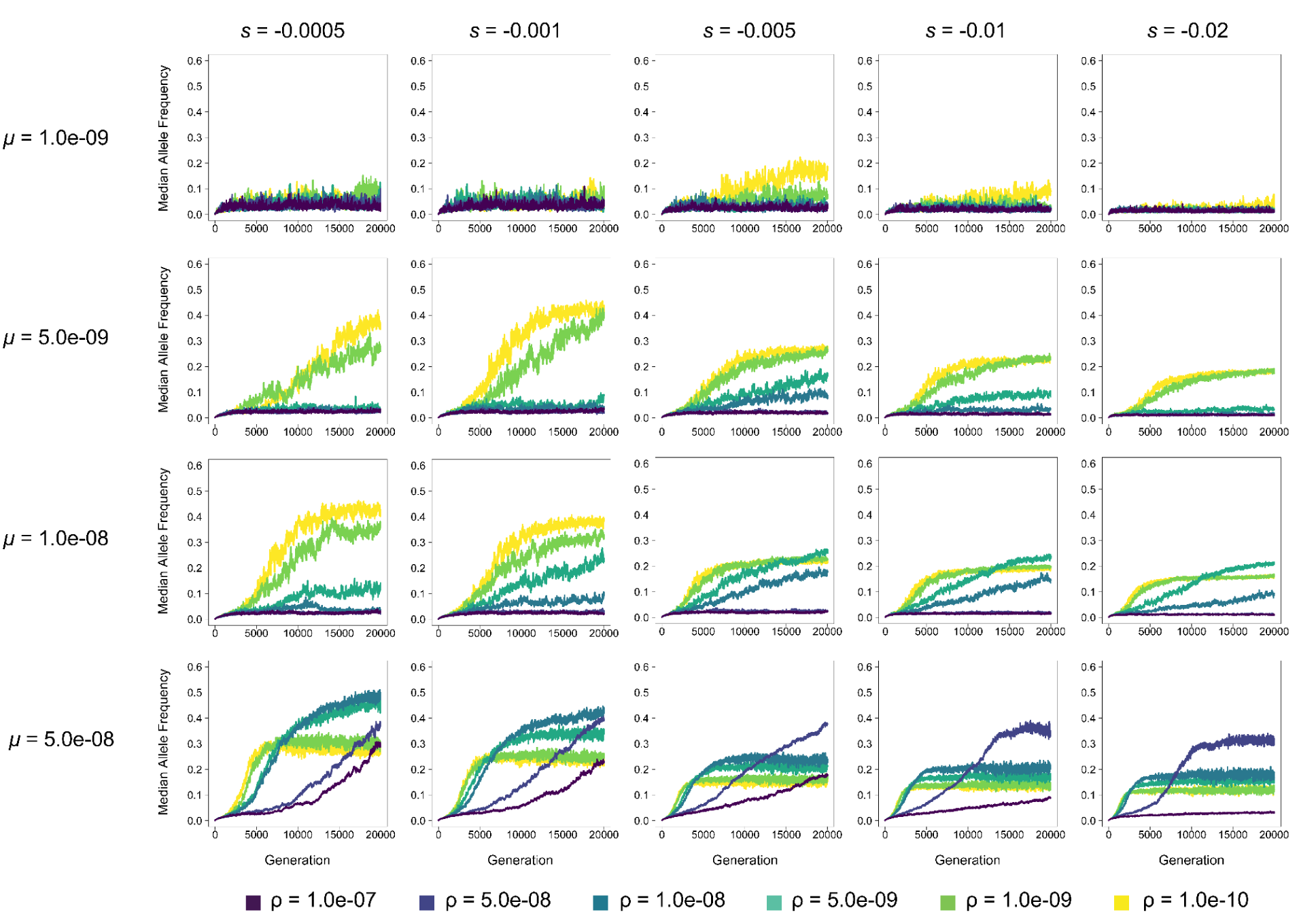
Median allele frequency of diploids in each generation at varying mutation rates (*μ*) and selection coefficients (*s*) for *N* = 500. Each line represents the average across 10 replicates. Note: y-axis is identical across subfigures.

**Supplemental Figure 10.**
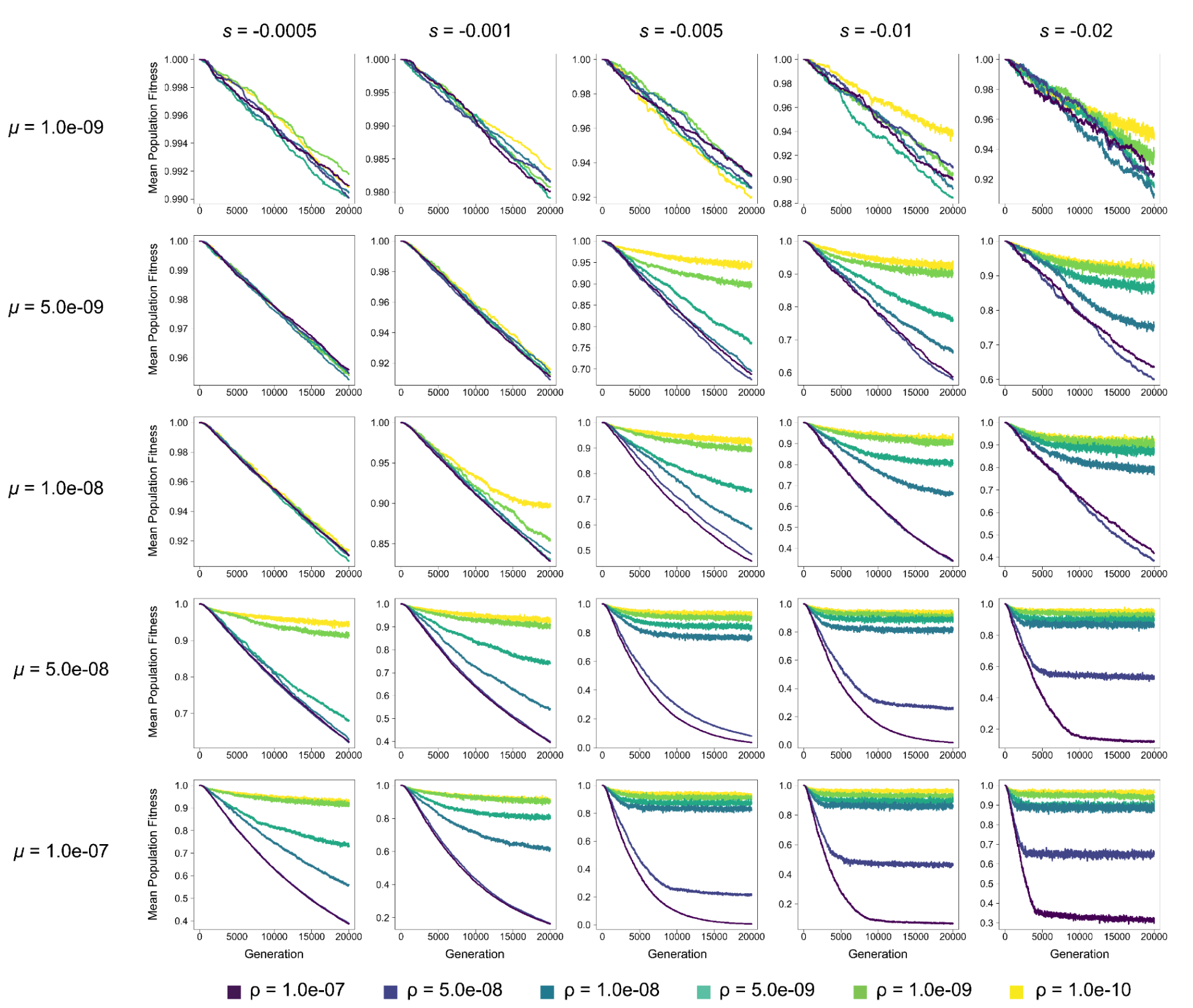
Mean population fitness of autotetraploids in each generation at varying mutation rates (*μ*) and selection coefficients (*s*) for *N* = 100. Each line represents the average across 10 replicates. Note: y-axis varies across subfigures.

**Supplemental Figure 11.**
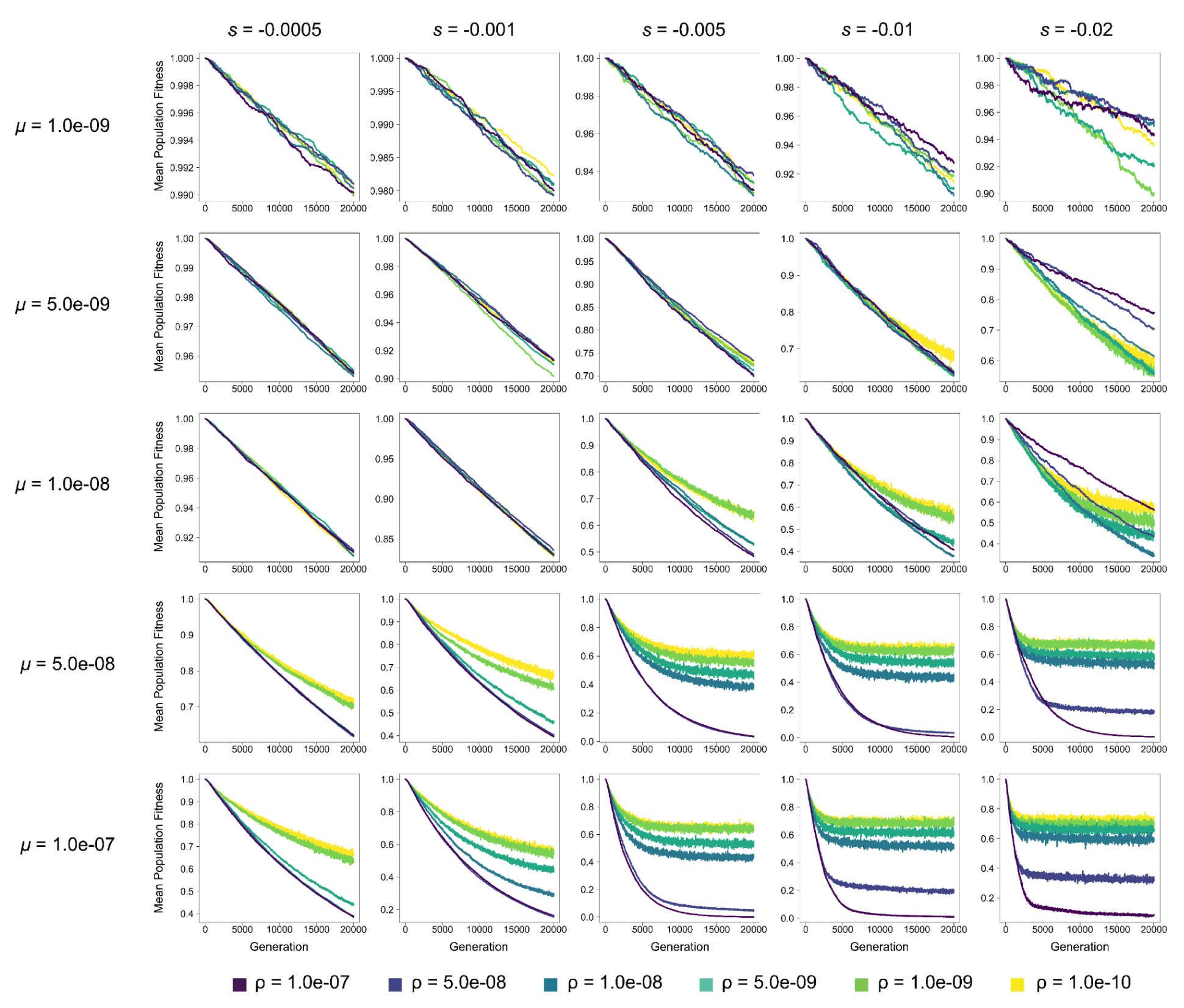
Mean population fitness of diploids in each generation at varying mutation rates (*μ*) and selection coefficients (*s*) for *N* = 100. Each line represents the average across 10 replicates. Note: y-axis varies across subfigures.

**Supplemental Figure 12.**
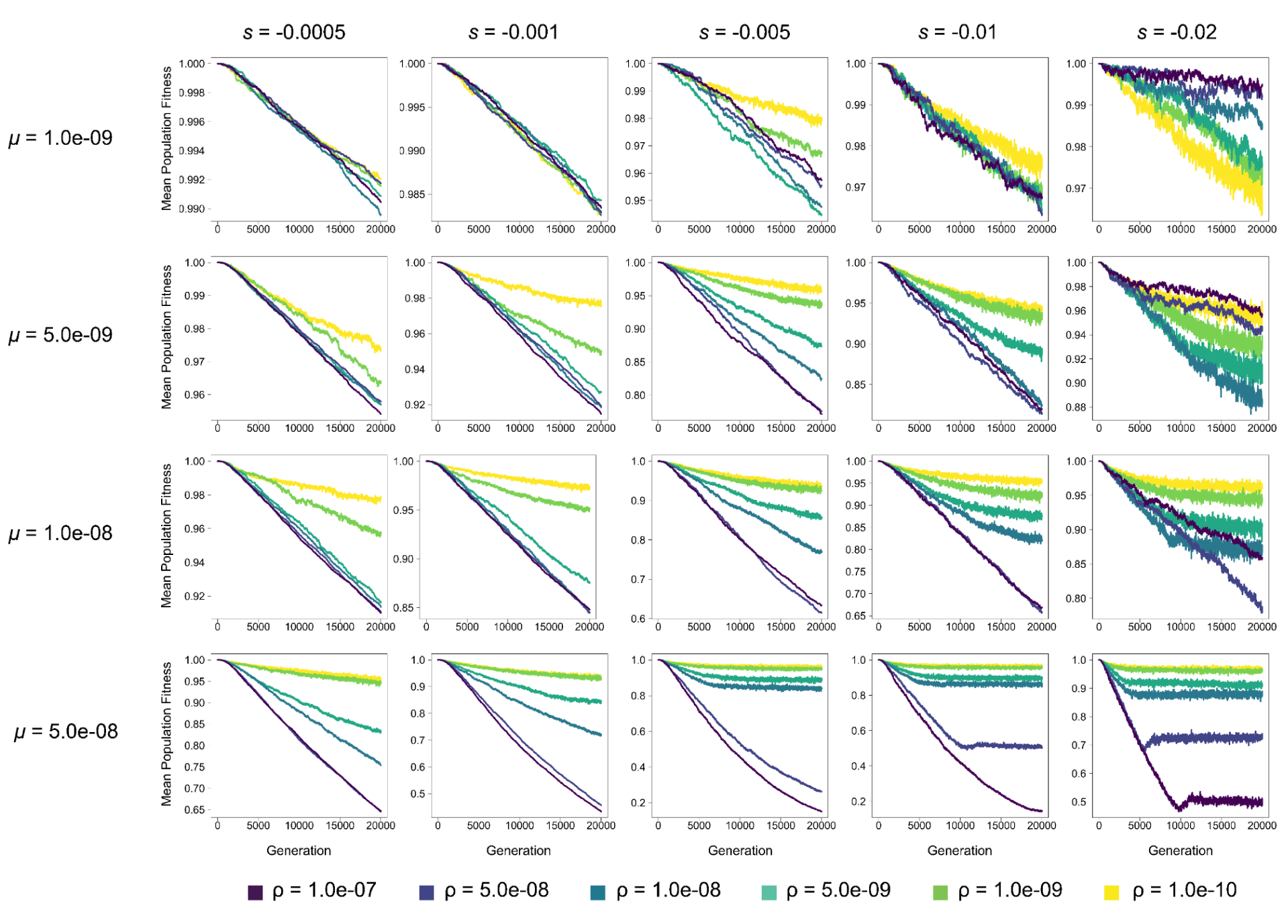
Mean population fitness of autopolyploids in each generation at varying mutation rates (*μ*) and selection coefficients (*s*) for *N* = 200. Each line represents the average across 10 replicates. Note: y-axis varies across subfigures.

**Supplemental Figure 13.**
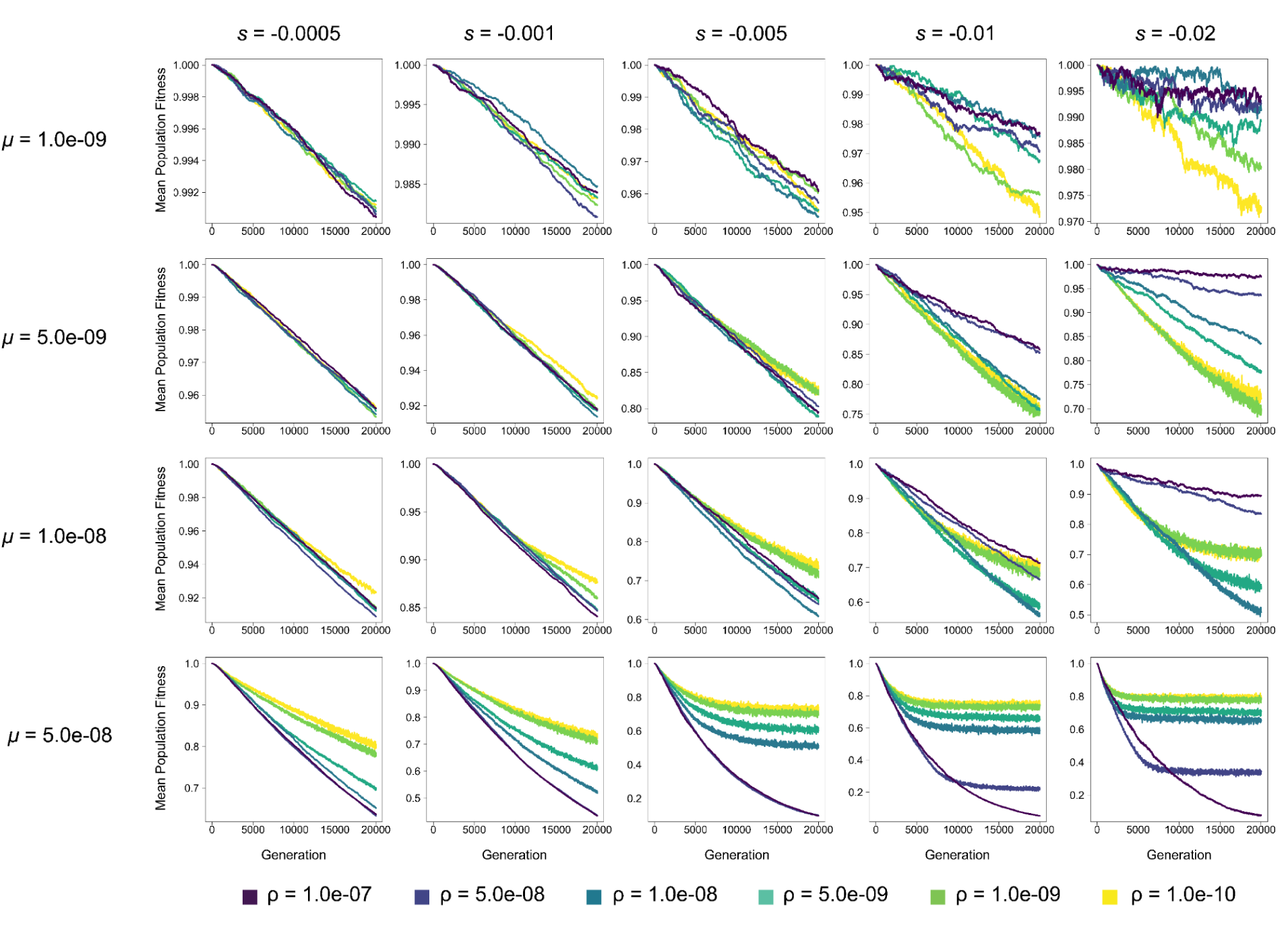
Mean population fitness of diploids in each generation at varying mutation rates (*μ*) and selection coefficients (*s*) for *N* = 200. Each line represents the average across 10 replicates. Note: y-axis varies across subfigures.

**Supplemental Figure 14.**
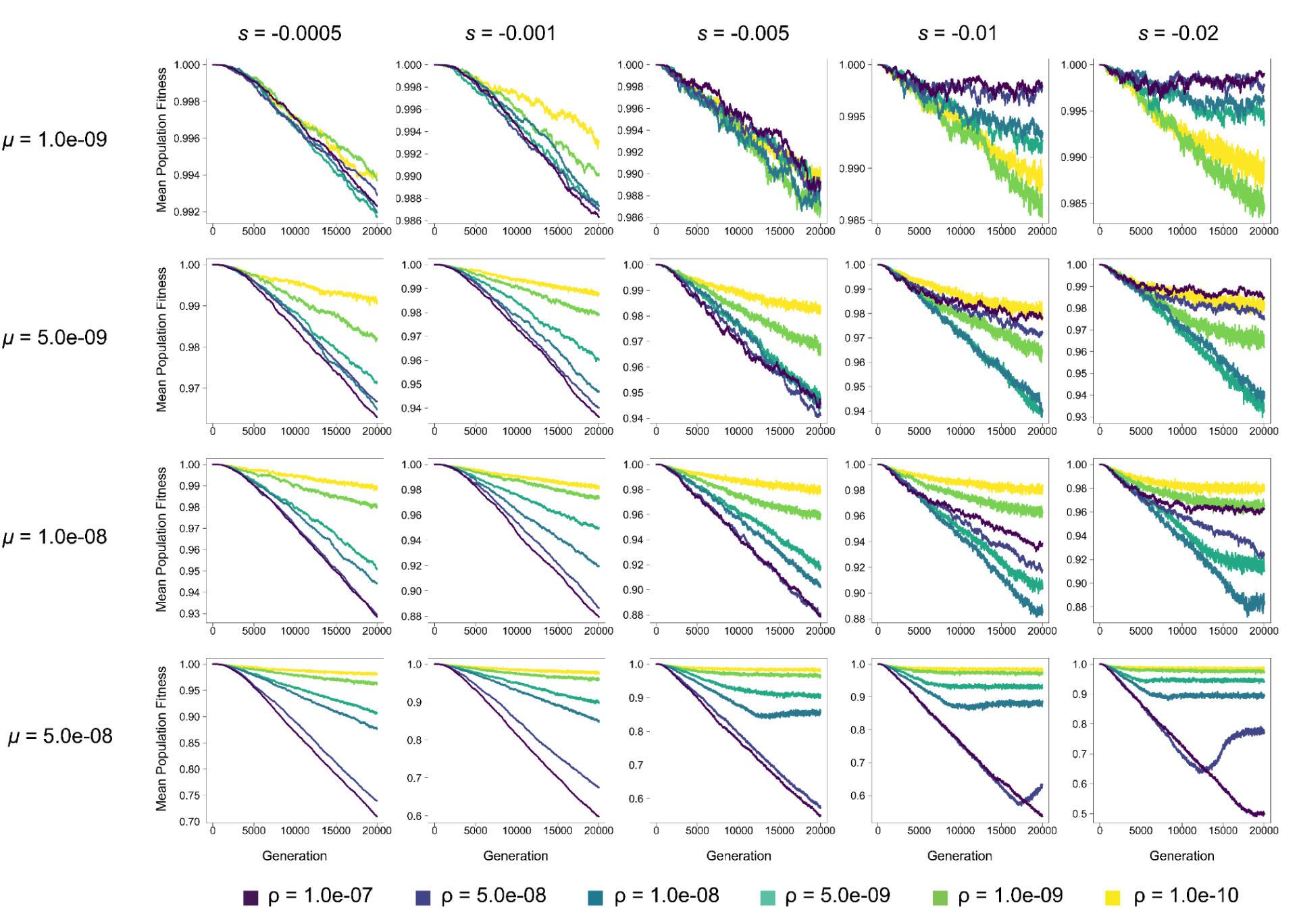
Mean population fitness of autopolyploids in each generation at varying mutation rates (*μ*) and selection coefficients (*s*) for *N* = 500. Each line represents the average across 10 replicates. Note: y-axis varies across subfigures.

**Supplemental Figure 15.**
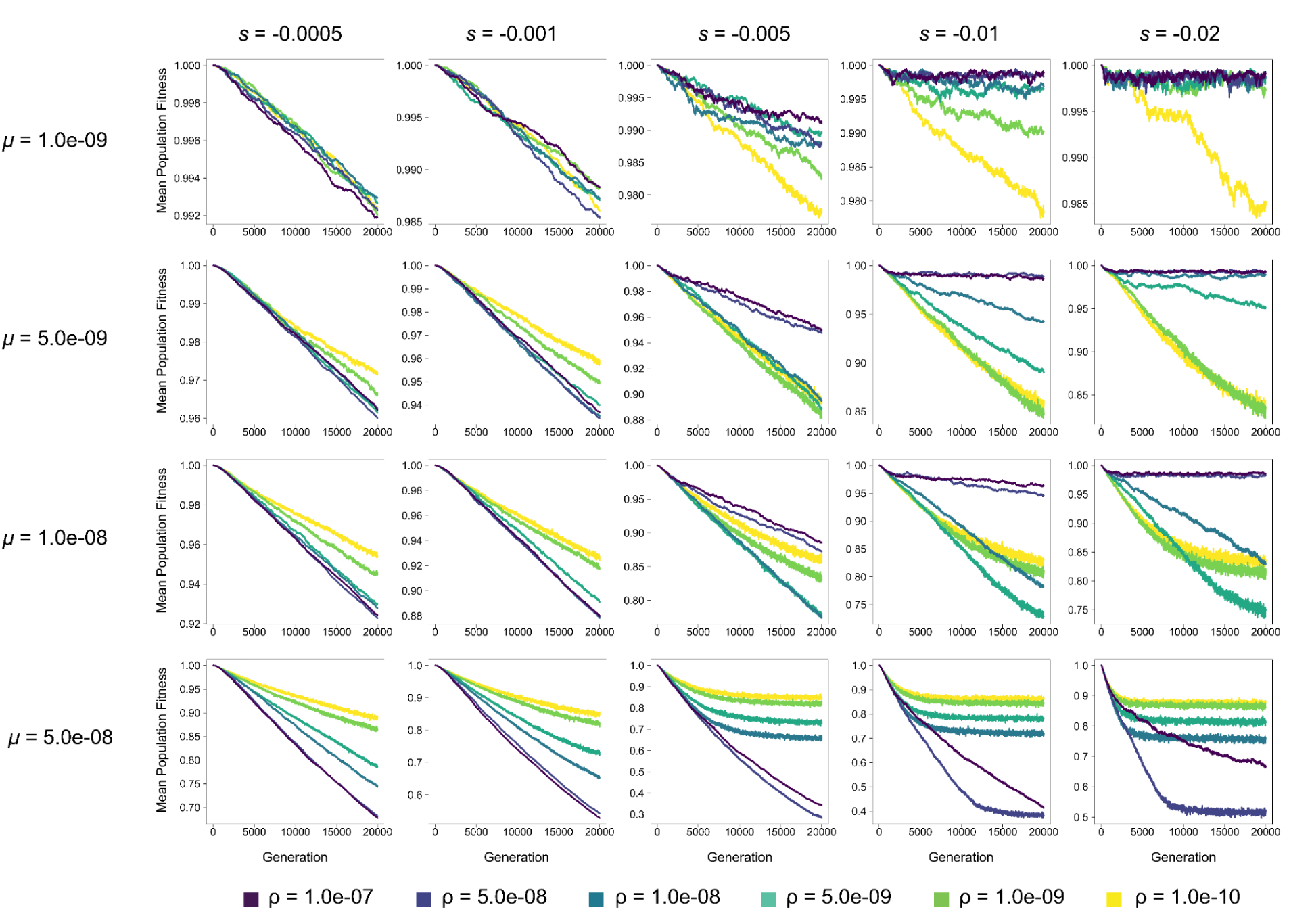
Mean population fitness of diploids in each generation at varying mutation rates (*μ*) and selection coefficients (*s*) for *N* = 500. Each line represents the average across 10 replicates. Note: y-axis varies across subfigures.

**Supplemental Figure 16.**
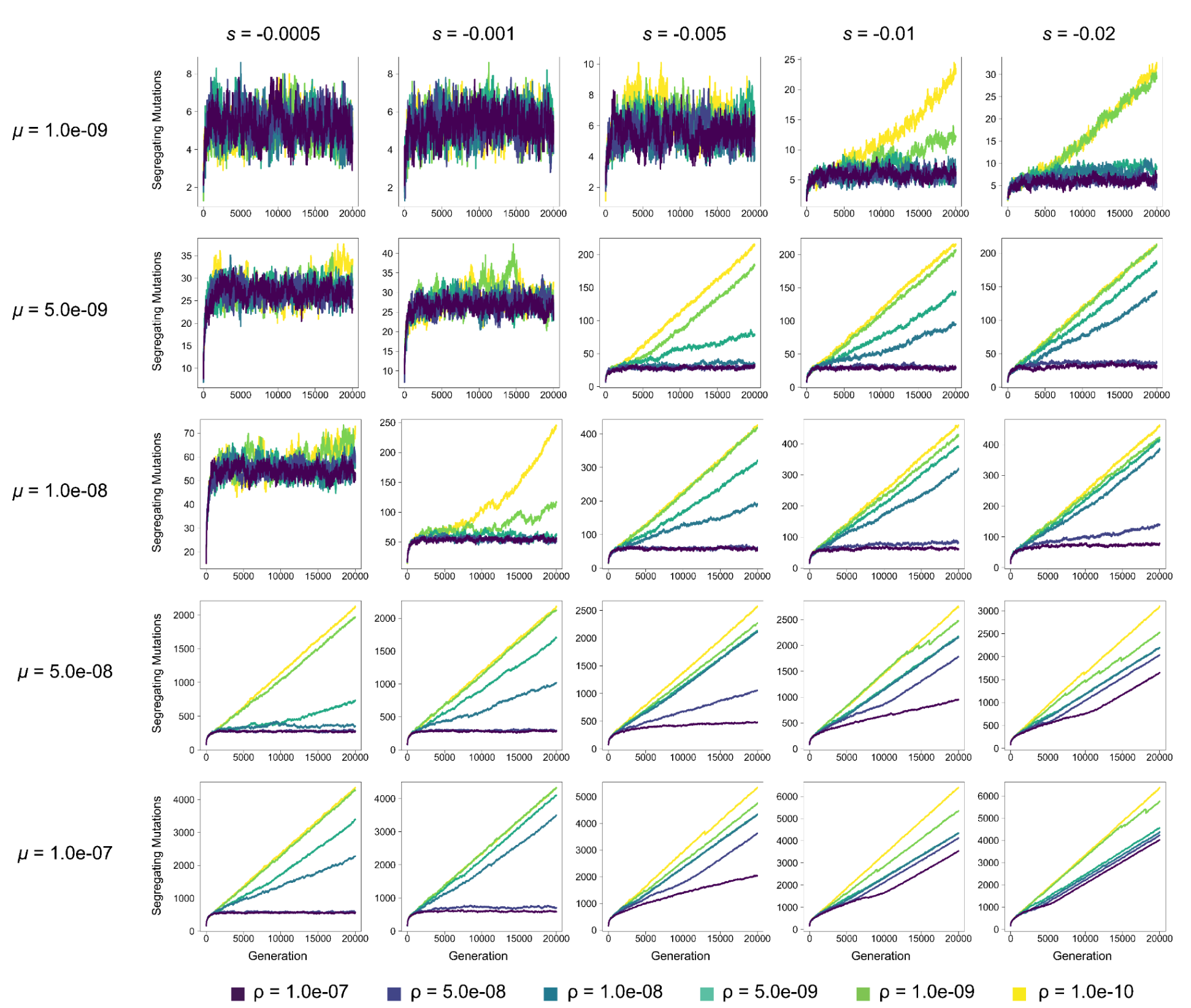
The number of segregating mutations in autotetraploid populations in each generation at varying mutation rates (*μ*) and selection coefficients (*s*) for *N* = 100. Each line represents the average across 10 replicates. Note: y-axis varies across subfigures.

**Supplemental Figure 17.**
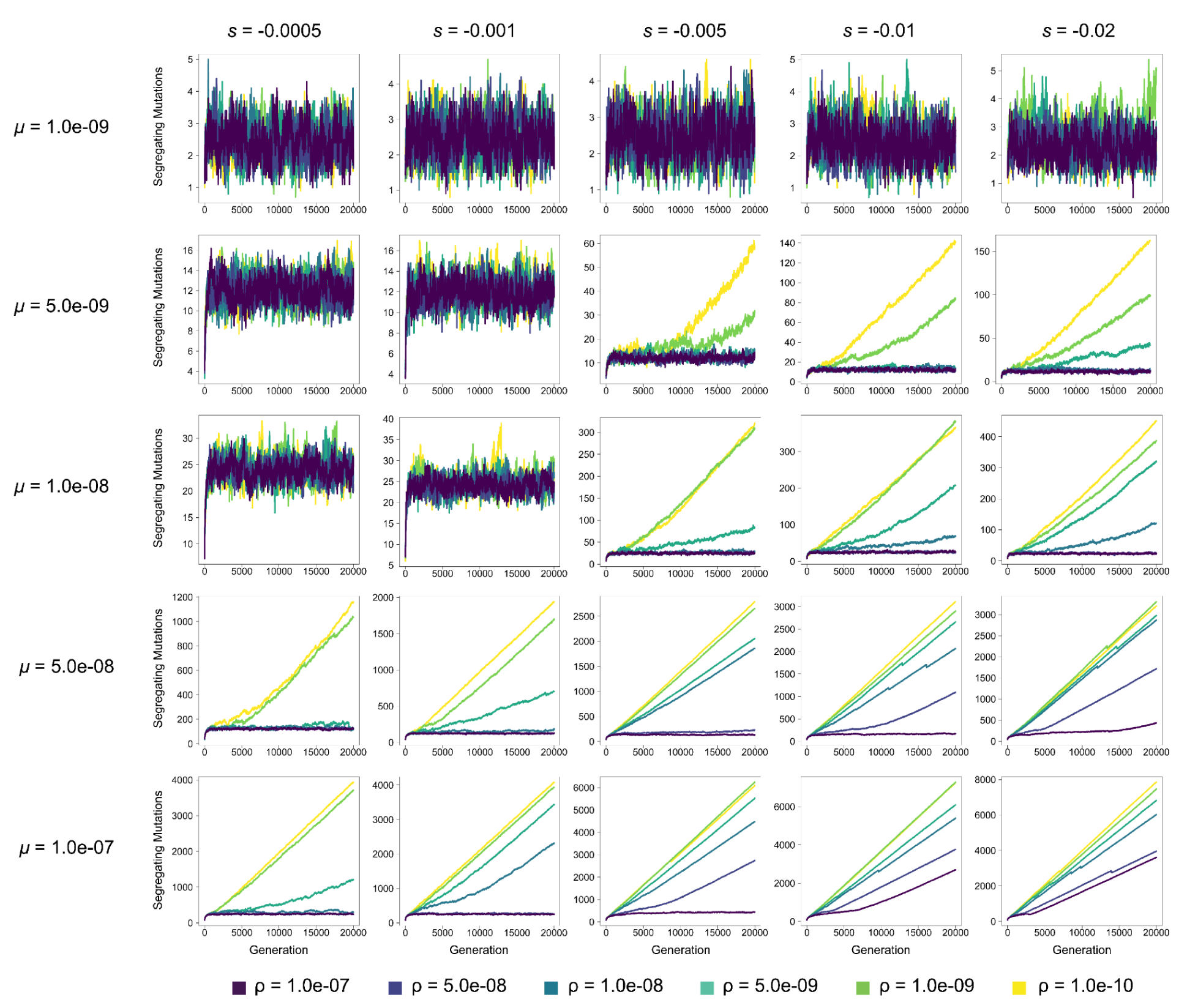
The number of segregating mutations in diploid populations in each generation at varying mutation rates (*μ*) and selection coefficients (*s*) for *N* = 100.Each line represents the average across 10 replicates. Note: y-axis varies across subfigures.

**Supplemental Figure 18.**
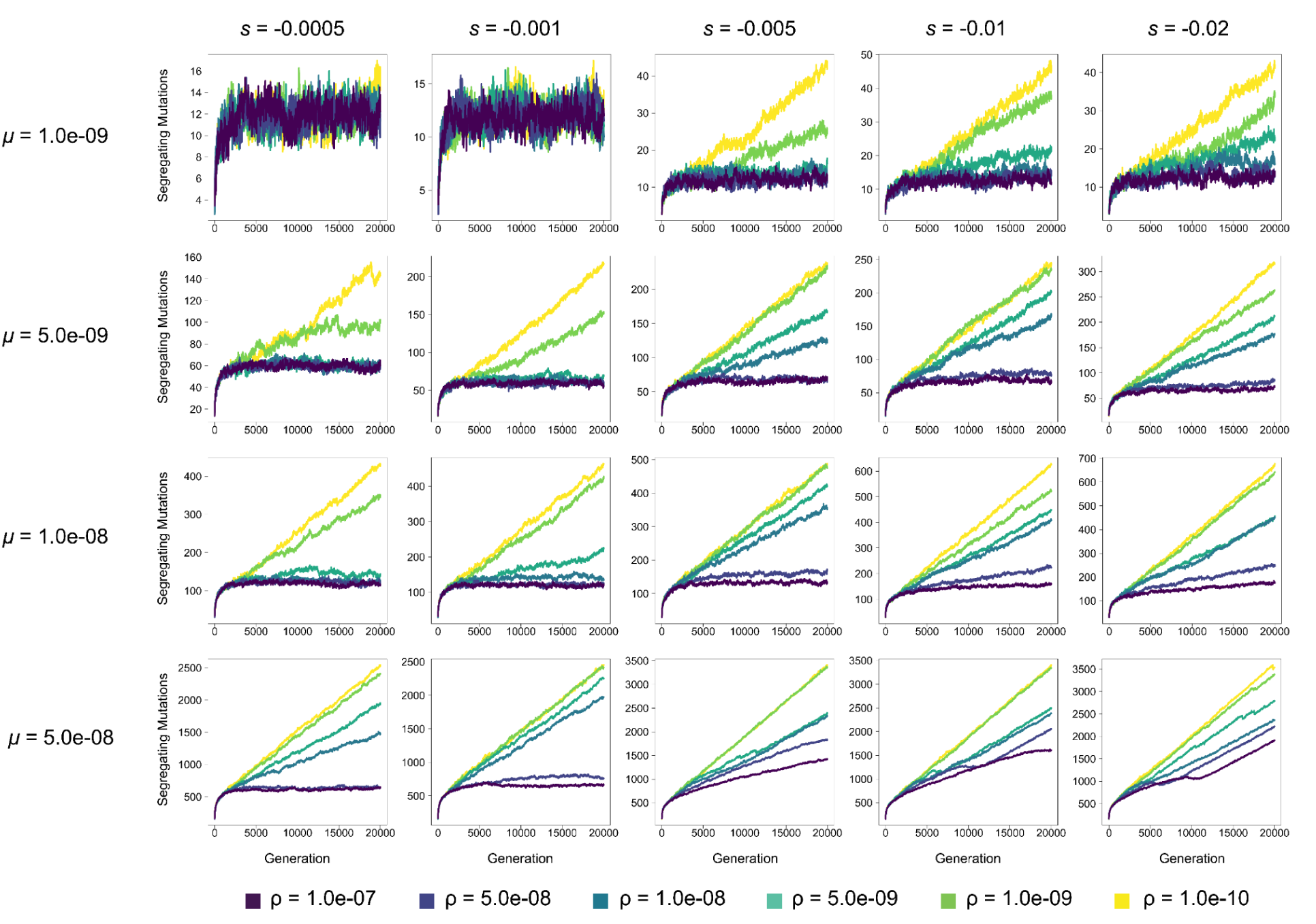
The number of segregating mutations in autotetraploid populations in each generation at varying mutation rates (*μ*) and selection coefficients (*s*) for *N* = 200. Each line represents the average across 10 replicates. Note: y-axis varies across subfigures.

**Supplemental Figure 19.**
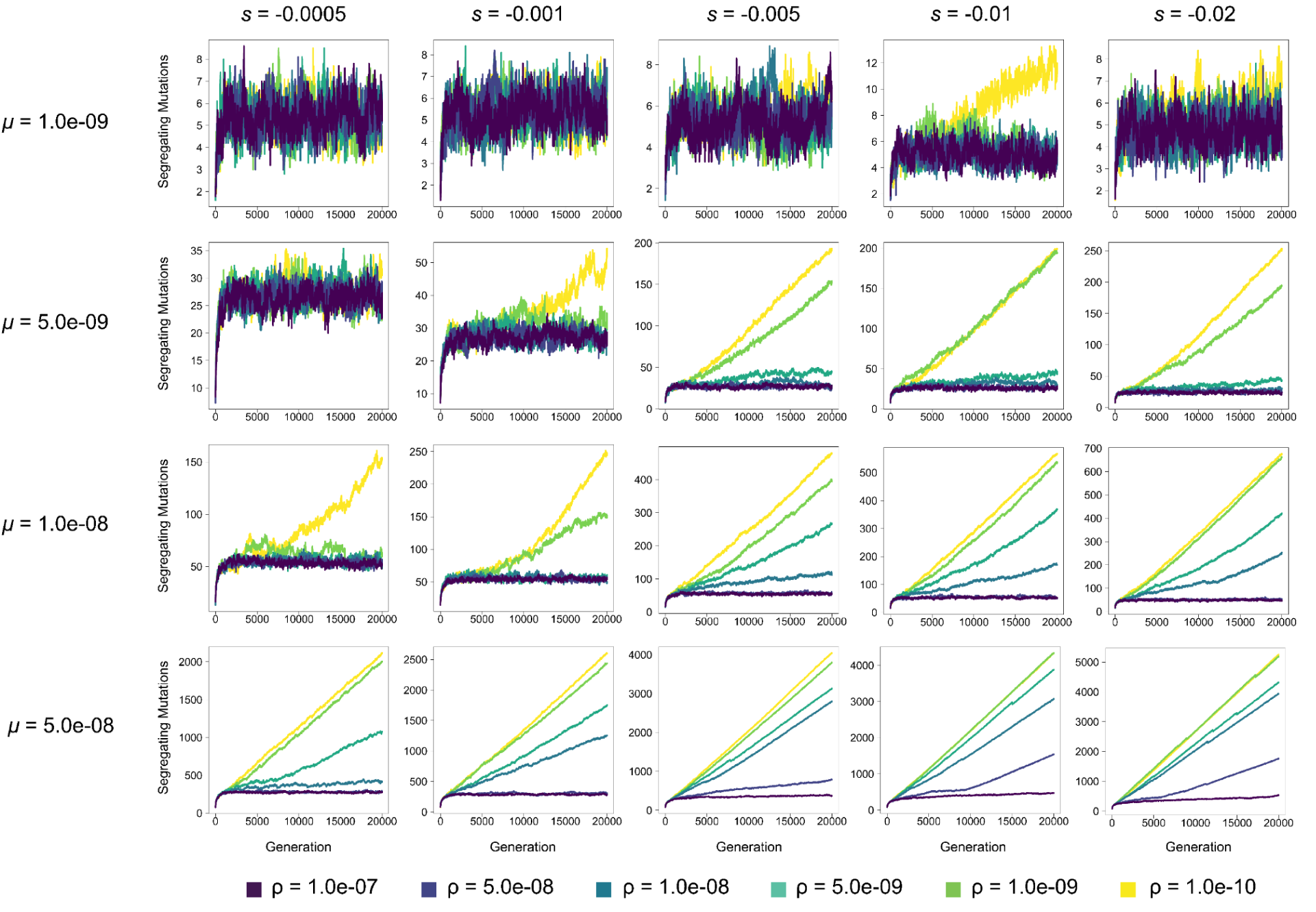
The number of segregating mutations in diploid populations in each generation at varying mutation rates (*μ*) and selection coefficients (*s*) for *N* = 200. Each line represents the average across 10 replicates. Note: y-axis varies across subfigures.

**Supplemental Figure 20.**
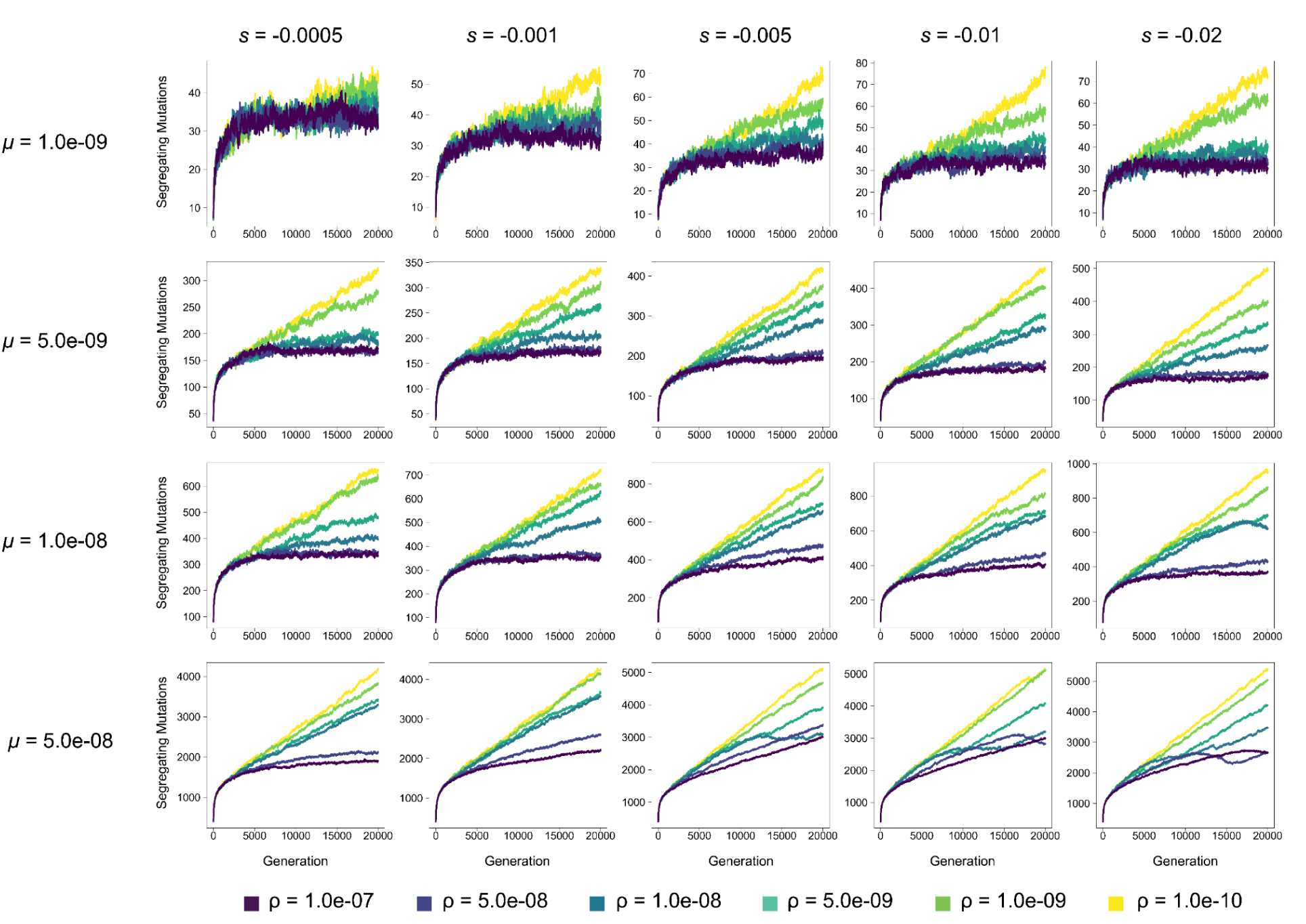
The number of segregating mutations in autotetraploid populations in each generation at varying mutation rates (*μ*) and selection coefficients (*s*) for *N* = 500. Each line represents the average across 10 replicates. Note: y-axis varies across subfigures.

**Supplemental Figure 21.**
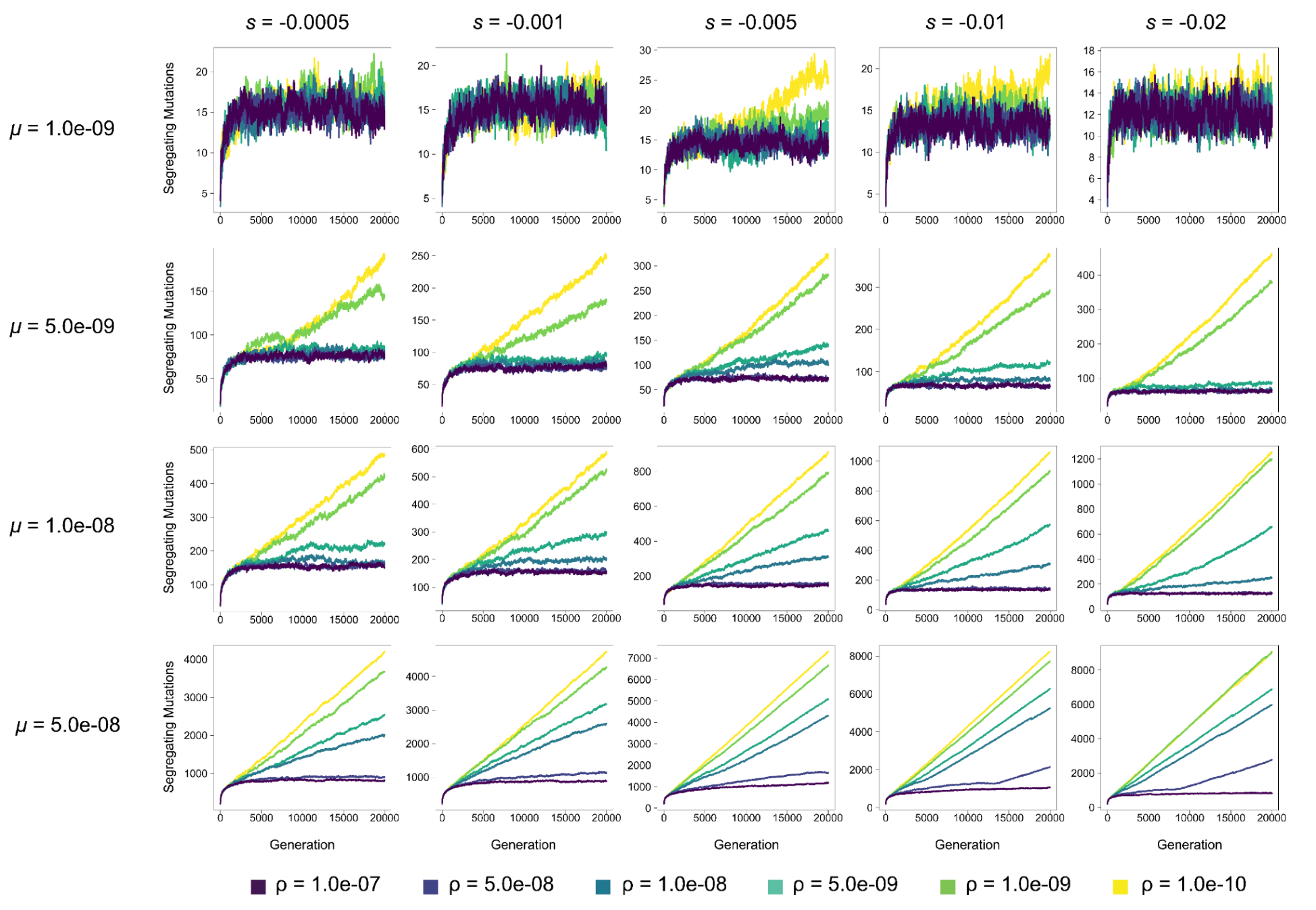
The number of segregating mutations in diploid populations in each generation at varying mutation rates (*μ*) and selection coefficients (*s*) for *N* = 500. Each line represents the average across 10 replicates. Note: y-axis varies across subfigures.

**Supplemental Figure 22.**
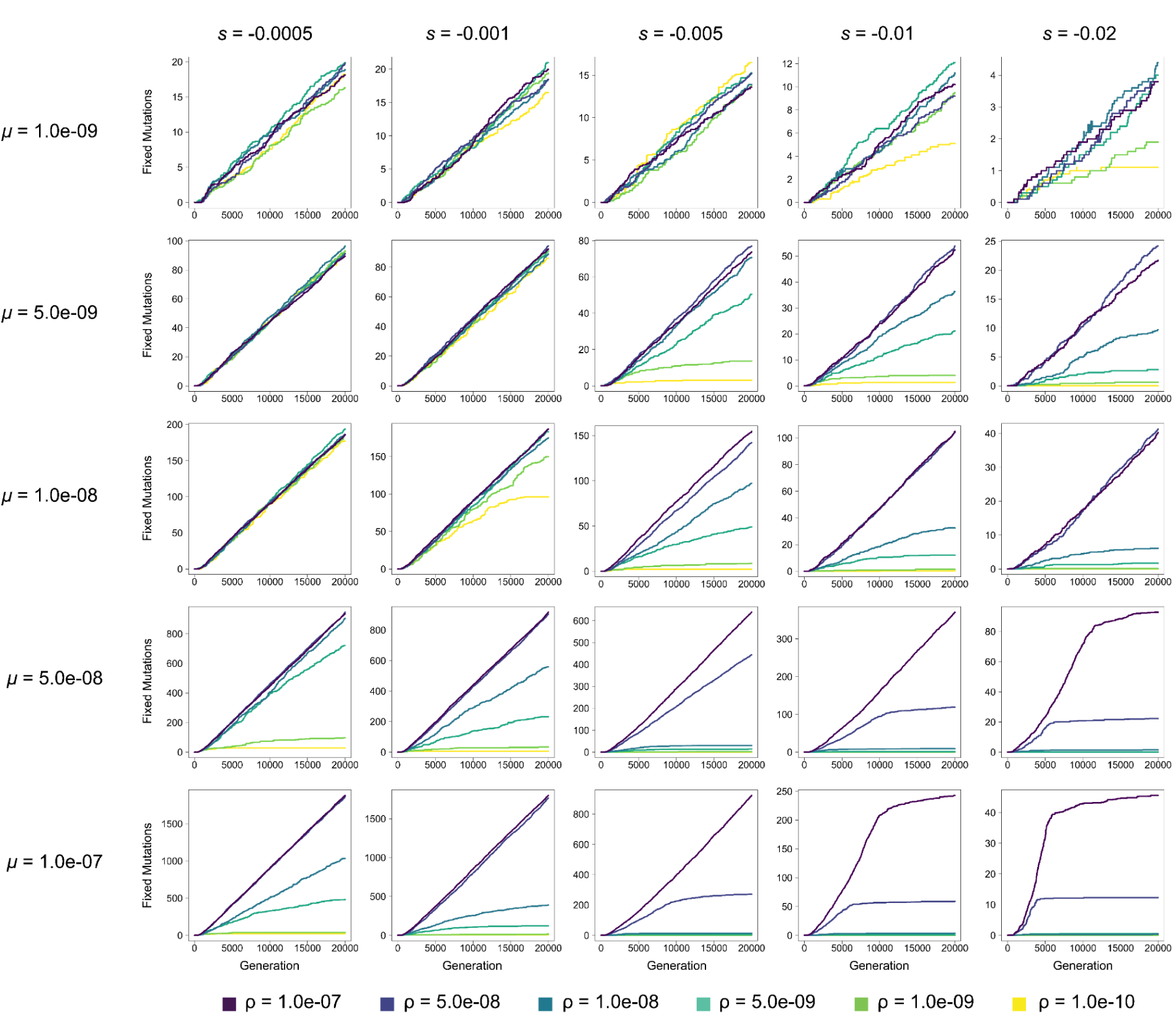
The number of fixed mutations in autotetraploid populations in each generation at varying mutation rates (*μ*) and selection coefficients (*s*) for *N* = 100. Each line represents the average across 10 replicates. Note: y-axis varies across subfigures.

**Supplemental Figure 23.**
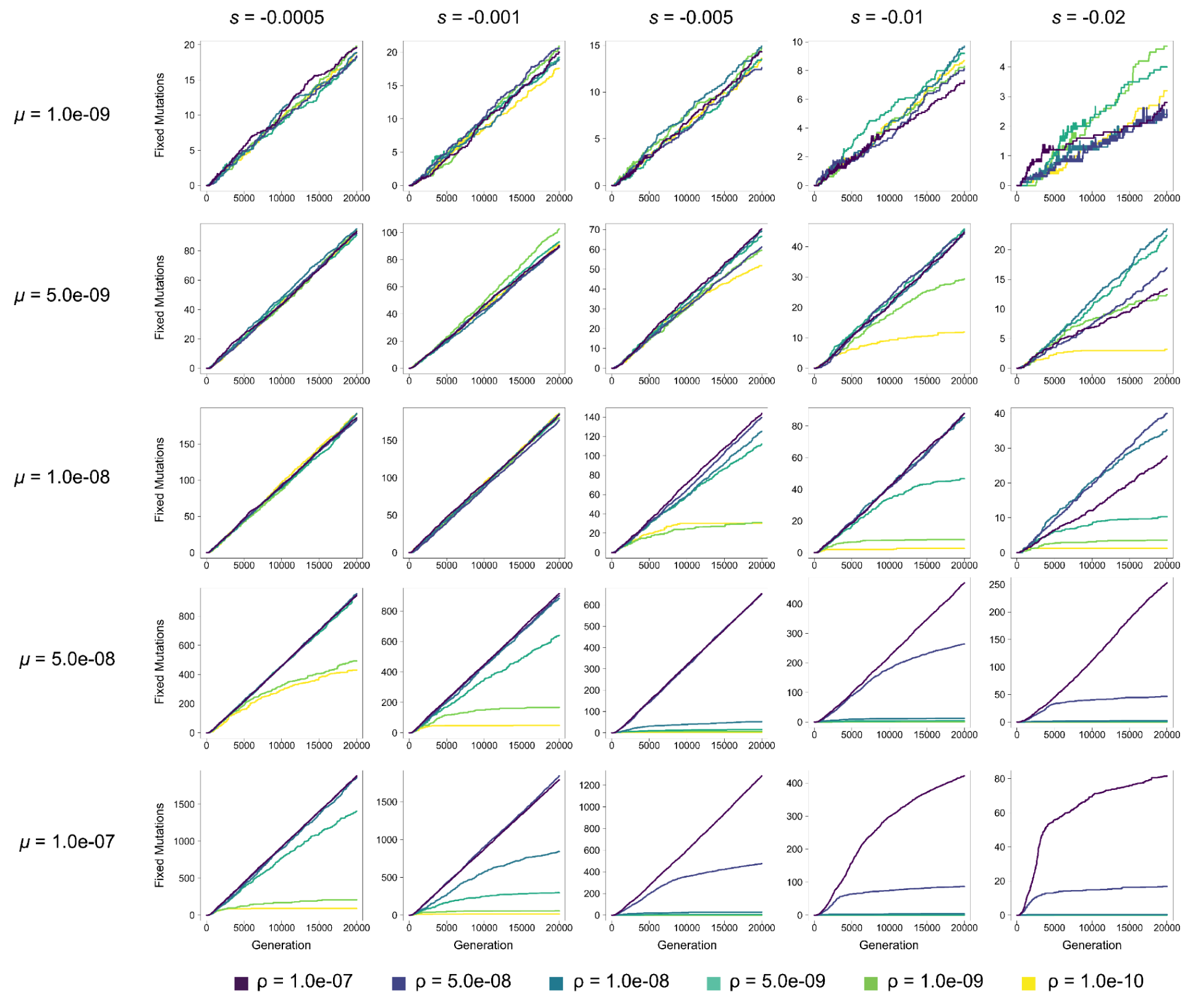
The number of fixed mutations in diploid populations in each generation at varying mutation rates (*μ*) and selection coefficients (*s*) for *N* = 100. Each line represents the average across 10 replicates. Note: y-axis varies across subfigures.

**Supplemental Figure 24.**
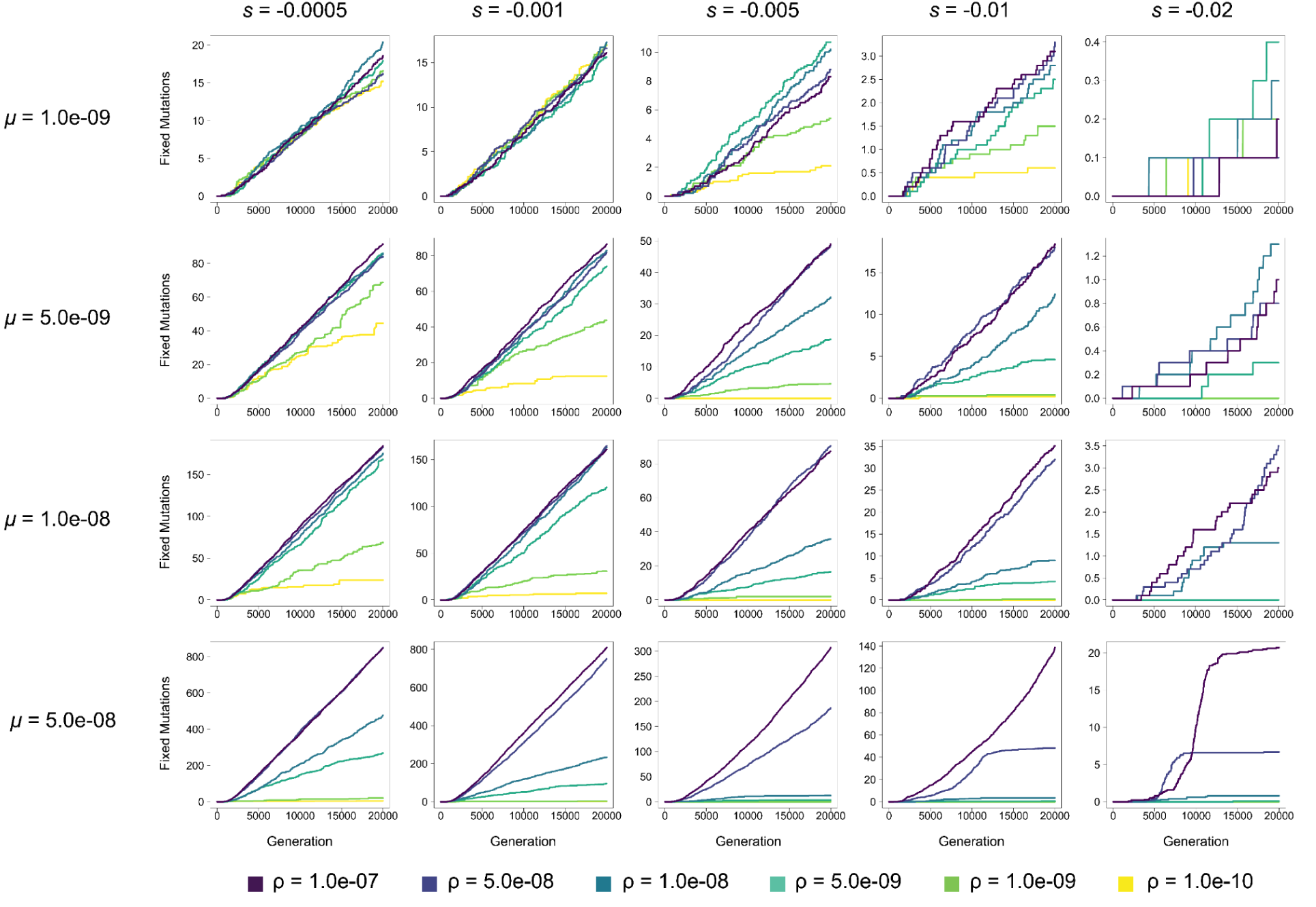
The number of fixed mutations in autotetraploid populations in each generation at varying mutation rates (*μ*) and selection coefficients (*s*) for *N* = 200. Each line represents the average across 10 replicates. Note: y-axis varies across subfigures.

**Supplemental Figure 25.**
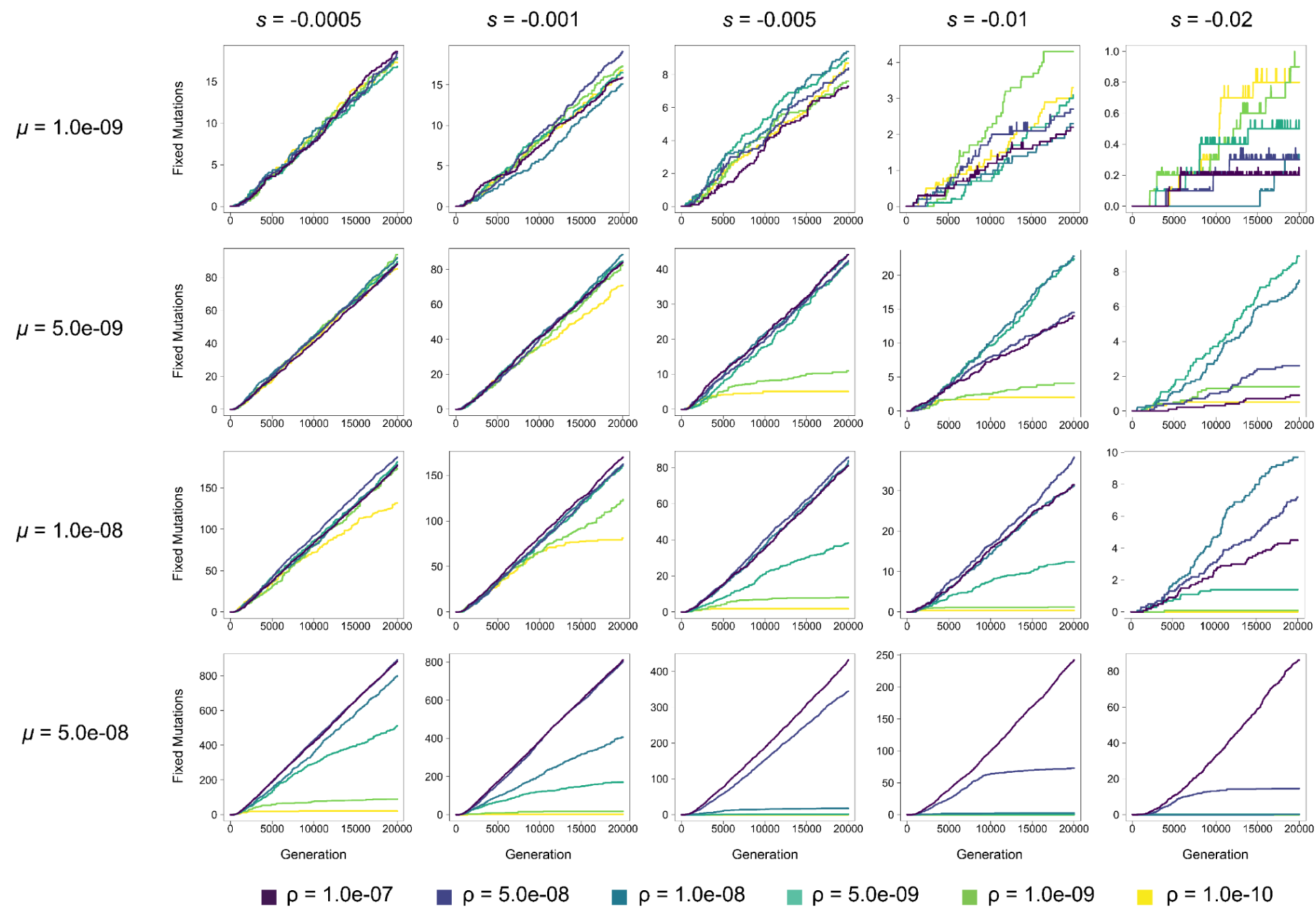
The number of fixed mutations in diploid populations in each generation at varying mutation rates (*μ*) and selection coefficients (*s*) for *N* = 200. Each line represents the average across 10 replicates. Note: y-axis varies across subfigures.

**Supplemental Figure 26.**
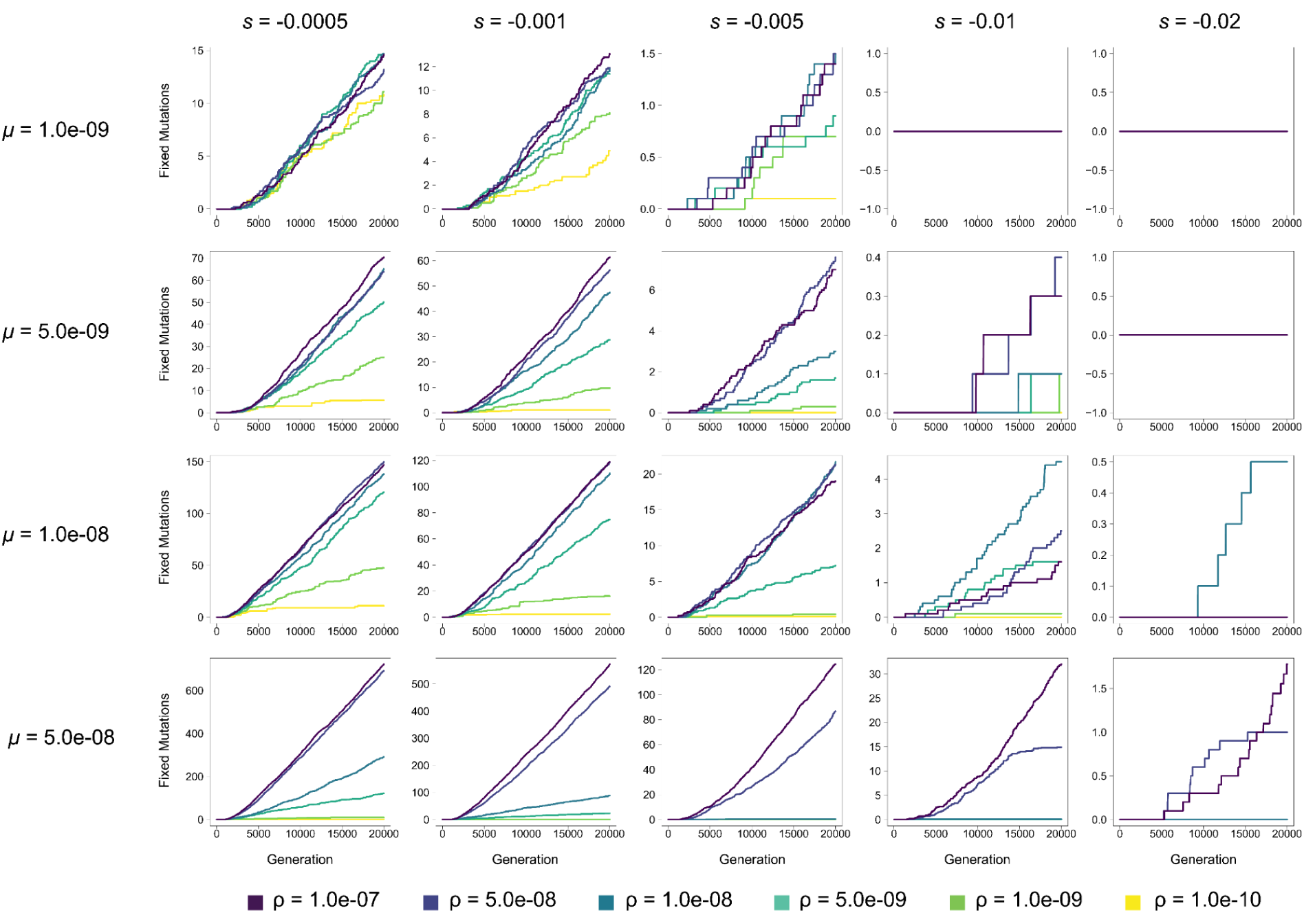
The number of fixed mutations in autotetraploid populations in each generation at varying mutation rates (*μ*) and selection coefficients (*s*) for *N* = 500. Each line represents the average across 10 replicates. Note: y-axis varies across subfigures.

**Supplemental Figure 27.**
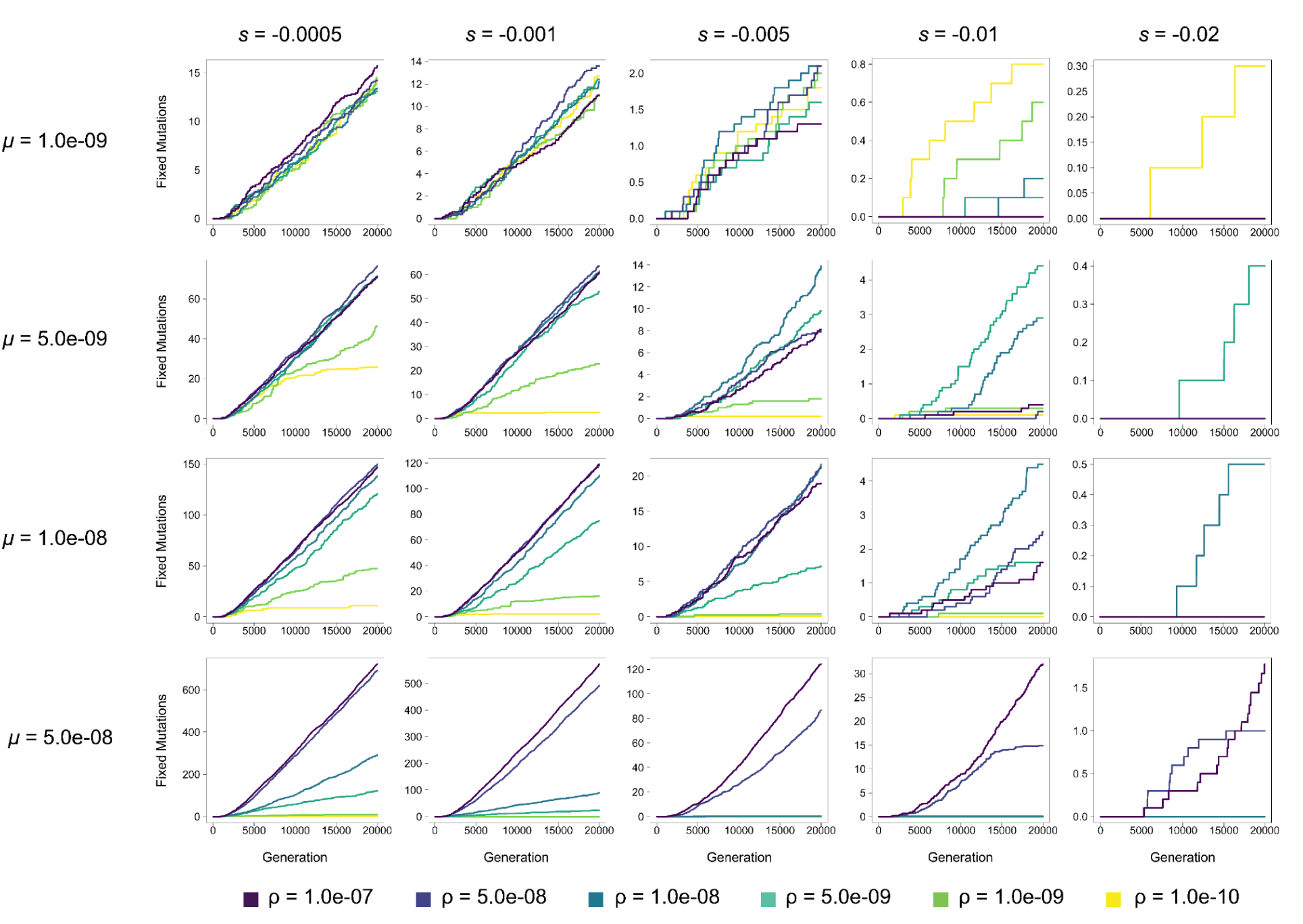
The number of fixed mutations in diploid populations in each generation at varying mutation rates (*μ*) and selection coefficients (*s*) for *N* = 500. Each line represents the average across 10 replicates. Note: y-axis varies across subfigures.

## Reference

Albertin W, Marullo P. 2012. Polyploidy in fungi: evolution after whole-genome duplication. Proc Biol Sci. 279(1738):2497–2509. doi:10.1098/rspb.2012.0434.

Amalraj VA, Balasundaram N. 2006. On the taxonomy of the members of ‘saccharum complex.’ Genet Resour Crop Evol. 53(1):35–41. doi:10.1007/s10722-004-0581-1.

Barker MS, Husband BC, Pires JC. 2016. Spreading Winge and flying high: The evolutionary importance of polyploidy after a century of study. Am J Bot. 103(7):1139–1145. doi:10.3732/ajb.1600272.

Barringer BC. 2007. Polyploidy and self-fertilization in flowering plants. Am J Bot. 94(9):1527–1533. doi:10.3732/ajb.94.9.1527.

Barton NH, Charlesworth B. 1998. Why sex and recombination? Science. 281(5385):1986–1990. doi:10.1126/science.281.5385.1986.

Barton NH. 1995. A general model for the evolution of recombination. Genet Res. 65(2):123–145. doi:10.1017/s0016672300033140.

Becher H, Jackson BC, Charlesworth B. 2020. Patterns of Genetic Variability in Genomic Regions with Low Rates of Recombination. Curr Biol. 30(1):94–100.e3. doi:10.1016/j.cub.2019.10.047.

Bierne N, Tsitrone A, David P. 2000. An inbreeding model of associative overdominance during a population bottleneck. Genetics. 155(4):1981–1990. doi:10.1093/genetics/155.4.1981.

Bomblies K, Jones G, Franklin C, Zickler D, Kleckner N. 2016. The challenge of evolving stable polyploidy: could an increase in “crossover interference distance” play a central role? Chromosoma. 125(2):287–300. doi:10.1007/s00412-015-0571-4.

Booker WW, Schrider DR. 2025. The genetic consequences of range expansion and its influence on diploidization in polyploids. The American Naturalist. 205(2):203–223. doi:10.1086/733334.

Brazier T, Glémin S. 2022. Diversity and determinants of recombination landscapes in flowering plants. PLoS Genet. 18(8):e1010141. doi:10.1371/journal.pgen.1010141.

Breuert S, Allers T, Spohn G, Soppa J. 2006. Regulated polyploidy in halophilic archaea. PLoS ONE. 1(1):e92. doi:10.1371/journal.pone.0000092.

Charlesworth B, Charlesworth D. 1975. An experimental on recombination load in Drosophila melanogaster. Genet Res. 25(3):267–274. doi:10.1017/s001667230001569x.

Charlesworth B, Jensen JD. 2021. Effects of selection at linked sites on patterns of genetic variability. Annu Rev Ecol Evol Syst. 52(1):177–197. doi:10.1146/annurev-ecolsys-010621-044528.

Charlesworth D. 1991. The apparent selection on neutral marker loci in partially inbreeding populations. Genet Res. 57(2):159–175. doi:10.1017/S0016672300029244.

Crow JF, Kimura M. 1965. Evolution in sexual and asexual populations. Am Nat. 99(909):439–450. doi:10.1086/282389.

Douches DS, Maas D, Jastrzebski K, Chase RW. 1996. Assessment of Potato Breeding Progress in the USA over the Last Century. Crop Sci. 36(6):1544–1552. doi:10.2135/cropsci1996.0011183X003600060024x.

Felsenstein J. 1974. The evolutionary advantage of recombination. Genetics. 78(2):737–756. doi:10.1093/genetics/78.2.737.

Ferreira FM, Barbosa MHP, Castro RD, Peternelli LA, Cruz CD. 2005. Effects of inbreeding on the selection of sugar cane clones. CBAB. 5(2):174–182. doi:10.12702/1984-7033.v05n02a07.

Frydenberg O. 1963. Population studies of a lethal mutant in drosophila melanogaster. Hereditas. 50(1):89–116. doi:10.1111/j.1601-5223.1963.tb01896.x.

Gilbert KJ, Pouyet F, Excoffier L, Peischl S. 2020. Transition from Background Selection to Associative Overdominance Promotes Diversity in Regions of Low Recombination. Curr Biol. 30(1):101–107.e3. doi:10.1016/j.cub.2019.11.063.

Gregory TR, Mable BK. 2005. Polyploidy in Animals. In: The evolution of the genome. Elsevier. (Gregory T. Ryan, editor.). p. 427–517.

Haller BC, Messer PW. 2017. Slim 2: flexible, interactive forward genetic simulations. Mol Biol Evol. 34(1):230–240. doi:10.1093/molbev/msw211.

Haller BC, Messer PW. 2023. SLiM 4: Multispecies Eco-Evolutionary Modeling. Am Nat. 201(5):E127–E139. doi:10.1086/723601.

Healey AL, Garsmeur O, Lovell JT, Shengquiang S, Sreedasyam A, Jenkins J, Plott CB, Piperidis N, Pompidor N, Llaca V, et al. 2024. The complex polyploid genome architecture of sugarcane. Nature. 628(8009):804–810. doi:10.1038/s41586-024-07231-4.

Hill WG, Robertson A. 1966. The effect of linkage on limits to artificial selection. Genet Res. 8(03):269–294. doi:10.1017/S0016672300010156.

Huber CD, Durvasula A, Hancock AM, Lohmueller KE. 2018. Gene expression drives the evolution of dominance. Nat Commun. 9(1):2750. doi:10.1038/s41467-018-05281-7.

Kyriazis CC, Wayne RK, Lohmueller KE. 2021. Strongly deleterious mutations are a primary determinant of extinction risk due to inbreeding depression. Evol Lett. 5(1):33–47. doi:10.1002/evl3.209.

Latter BD. 1998. Mutant alleles of small effect are primarily responsible for the loss of fitness with slow inbreeding in Drosophila melanogaster. Genetics. 148(3):1143–1158. doi:10.1093/genetics/148.3.1143.

Leary RF, Allendorf FW, Knudsen KL. 1984. Superior developmental stability of heterozygotes at enzyme loci in salmonid fishes. The American Naturalist. 124(4):540–551. doi:10.1086/284293.

van Lieshout N, van der Burgt A, de Vries ME, Ter Maat M, Eickholt D, Esselink D, van Kaauwen MPW, Kodde LP, Visser RGF, Lindhout P, et al. 2020. Solyntus, the New Highly Contiguous Reference Genome for Potato (Solanum tuberosum). G3 (Bethesda). 10(10):3489–3495. doi:10.1534/g3.120.401550.

Lindhout P, Meijer D, Schotte T, Hutten RCB, Visser RGF, van Eck HJ. 2011. Towards F1 hybrid seed potato breeding. Potato Res. 54(4):301–312. doi:10.1007/s11540-011-9196-z.

Li Z, McKibben MTW, Finch GS, Blischak PD, Sutherland BL, Barker MS. 2021. Patterns and processes of diploidization in land plants. Annu Rev Plant Biol. 72:387–410. doi:10.1146/annurev-arplant-050718-100344.

Li Z, Tiley GP, Galuska SR, Reardon CR, Kidder TI, Rundell RJ, Barker MS. 2018. Multiple large-scale gene and genome duplications during the evolution of hexapods. Proc Natl Acad Sci USA. 115(18):4713–4718. doi:10.1073/pnas.1710791115.

Morgan C, White MA, Franklin FCH, Zickler D, Kleckner N, Bomblies K. 2021. Evolution of crossover interference enables stable autopolyploidy by ensuring pairwise partner connections in Arabidopsis arenosa. Curr Biol. 31(21):4713–4726.e4. doi:10.1016/j.cub.2021.08.028.

Morrison JW, Rajhathy T. 1960. Chromosome behaviour in autotetraploid cereals and grasses. Chromosoma. 11(1):297–309. doi:10.1007/BF00328656.

Muller HJ. 1916. The Mechanism of Crossing-Over. Am Nat. 50(592):193–221. doi:10.1086/279534.

Muller HJ. 1964. The relation of recombination to mutational advance. Mutat Res. 106:2–9. doi:10.1016/0027-5107(64)90047-8.

Mulligan GA. 1967. Diploid and autotetraploid physaria vitulifera (cruciferae). Can J Bot. 45(2):183–188. doi:10.1139/b67-014.

Neiman M, Sharbel TF, Schwander T. 2014. Genetic causes of transitions from sexual reproduction to asexuality in plants and animals. J Evol Biol. 27(7):1346–1359. doi:10.1111/jeb.12357.

Ohta T, Kimura M. 1970. Development of associative overdominance through linkage disequilibrium in finite populations. Genet Res. 16(2):165–177. doi:10.1017/s0016672300002391.

Ohta T. 1971. Associative overdominance caused by linked detrimental mutations. Genet Res. 18(3):277–286. doi:10.1017/S0016672300012684.

Ohta T. 1973. Effect of linkage on behavior of mutant genes in finite populations. Theor Popul Biol. 4(2):145–162. doi:10.1016/0040-5809(73)90025-7.

One Thousand Plant Transcriptomes Initiative. 2019. One thousand plant transcriptomes and the phylogenomics of green plants. Nature. 574(7780):679–685. doi:10.1038/s41586-019-1693-2.

Otto SP, Whitton J. 2000. Polyploid Incidence and Evolution. Annual Review of Genetics. 34:401–437.

Pálsson S. 2001. The effects of deleterious mutations in cyclically parthenogenetic organisms. J Theor Biol. 208(2):201–214. doi:10.1006/jtbi.2000.2211.

Pálsson S. 2002. Selection on a modifier of recombination rate due to linked deleterious mutations. J Hered. 93(1):22–26. doi:10.1093/jhered/93.1.22.

Pamilo P, Pálsson S. 1998. Associative overdominance, heterozygosity and fitness. Heredity. 81 (Pt 4):381–389. doi:10.1046/j.1365-2540.1998.00395.x.

Raboin L-M, Pauquet J, Butterfield M, D’Hont A, Glaszmann J-C. 2008. Analysis of genome-wide linkage disequilibrium in the highly polyploid sugarcane. Theor Appl Genet. 116(5):701–714. doi:10.1007/s00122-007-0703-1.

Reddi VR. 1970. Pachytene pairing and the nature of polyploidy in Sorghum arundinaceum. Caryologia. 23(3):295–302.

Renny-Byfield S, Wendel JF. 2014. Doubling down on genomes: polyploidy and crop plants. Am J Bot. 101(10):1711–1725. doi:10.3732/ajb.1400119.

Ritz KR, Noor MAF, Singh ND. 2017. Variation in recombination rate: adaptive or not? Trends Genet. 33(5):364–374. doi:10.1016/j.tig.2017.03.003.

Roessler K, Muyle A, Diez CM, Gaut GRJ, Bousios A, Stitzer MC, Seymour DK, Doebley JF, Liu Q, Gaut BS. 2019. The genome-wide dynamics of purging during selfing in maize. Nat Plants. 5(9):980–990. doi:10.1038/s41477-019-0508-7.

Rougemont Q, Leroy T, Rondeau EB, Koop B, Bernatchez L. 2023. Allele surfing causes maladaptation in a Pacific salmon of conservation concern. PLoS Genet. 19(9):e1010918. doi:10.1371/journal.pgen.1010918.

Salman-Minkov A, Sabath N, Mayrose I. 2016. Whole-genome duplication as a key factor in crop domestication. Nat Plants. 2:16115. doi:10.1038/nplants.2016.115.

Stapley J, Feulner PGD, Johnston SE, Santure AW, Smadja CM. 2017. Variation in recombination frequency and distribution across eukaryotes: patterns and processes. Philos Trans R Soc Lond B Biol Sci. 372(1736). doi:10.1098/rstb.2016.0455.

Stebbins GL. 1950. Variation and evolution in plants. Columbia University Press.

Stenberg P, Saura A. 2013. Meiosis and its deviations in polyploid animals. Cytogenet Genome Res. 140(2–4):185–203. doi:10.1159/000351731.

Sved JA. 1968. The stability of linked systems of loci with a small population size. Genetics. 59(4):543–563. doi:10.1093/genetics/59.4.543.

Sved JA. 1971. Linkage disequilibrium and homozygosity of chromosome segments in finite populations. Theor Popul Biol. 2(2):125–141. doi:10.1016/0040-5809(71)90011-6.

Sved JA. 1972. Heterosis at the level of the chromosome and at the level of the gene. Theor Popul Biol. 3(4):491–506. doi:10.1016/0040-5809(72)90019-6.

Van de Peer Y, Mizrachi E, Marchal K. 2017. The evolutionary significance of polyploidy. Nat Rev Genet. 18(7):411–424. doi:10.1038/nrg.2017.26.

Waller DM. 2021. Addressing Darwin’s dilemma: Can pseudo-overdominance explain persistent inbreeding depression and load? Evolution. 75(4):779–793. doi:10.1111/evo.14189.

Wang S, Zickler D, Kleckner N, Zhang L. 2015. Meiotic crossover patterns: obligatory crossover, interference and homeostasis in a single process. Cell Cycle. 14(3):305–314. doi:10.4161/15384101.2014.991185.

Wilfert L, Gadau J, Schmid-Hempel P. 2007. Variation in genomic recombination rates among animal taxa and the case of social insects. Heredity. 98(4):189–197. doi:10.1038/sj.hdy.6800950.

Wood TE, Takebayashi N, Barker MS, Mayrose I, Greenspoon PB, Rieseberg LH. 2009. The frequency of polyploid speciation in vascular plants. Proc Natl Acad Sci USA. 106(33):13875–13879. doi:10.1073/pnas.0811575106.

Wu J-H, Datson PM, Manako KI, Murray BG. 2014. Meiotic chromosome pairing behaviour of natural tetraploids and induced autotetraploids of Actinidia chinensis. Theor Appl Genet. 127(3):549–557. doi:10.1007/s00122-013-2238-y.

Wu Y, Li D, Hu Y, Li H, Ramstein GP, Zhou S, Zhang X, Bao Z, Zhang Y, Song B, et al. 2023. Phylogenomic discovery of deleterious mutations facilitates hybrid potato breeding. Cell. 186(11):2313–2328.e15. doi:10.1016/j.cell.2023.04.008.

Yadav S, Jackson P, Wei X, Ross EM, Aitken K, Deomano E, Atkin F, Hayes BJ, Voss-Fels KP. 2020. Accelerating genetic gain in sugarcane breeding using genomic selection. Agronomy. 10(4):585. doi:10.3390/agronomy10040585.

Yant L, Hollister JD, Wright KM, Arnold BJ, Higgins JD, Franklin FCH, Bomblies K. 2013. Meiotic adaptation to genome duplication in Arabidopsis arenosa. Curr Biol. 23(21):2151–2156. doi:10.1016/j.cub.2013.08.059.

Zhang C, Wang P, Tang D, Yang Z, Lu F, Qi J, Tawari NR, Shang Y, Li C, Huang S. 2019. The genetic basis of inbreeding depression in potato. Nat Genet. 51(3):374–378. doi:10.1038/s41588-018-0319-1.

Zhang K, Wang X, Cheng F. 2019. Plant polyploidy: origin, evolution, and its influence on crop domestication. Horticultural Plant Journal. 5(6):231–239. doi:10.1016/j.hpj.2019.11.003.

Zhan SH, Drori M, Goldberg EE, Otto SP, Mayrose I. 2016. Phylogenetic evidence for cladogenetic polyploidization in land plants. Am J Bot. 103(7):1252–1258. doi:10.3732/ajb.1600108.

Zhao L, Charlesworth B. 2016. Resolving the conflict between associative overdominance and background selection. Genetics. 203(3):1315–1334. doi:10.1534/genetics.116.188912.

